# The epigenomic landscape regulating organogenesis in human embryos linked to developmental disorders

**DOI:** 10.1101/691766

**Authors:** Dave T. Gerrard, Andrew A. Berry, Rachel E. Jennings, Matthew J Birket, Sarah J Withey, Patrick Short, Sandra Jiménez-Gancedo, Panos N Firbas, Ian Donaldson, Andrew D. Sharrocks, Karen Piper Hanley, Matthew E Hurles, José Luis Gomez-Skarmeta, Nicoletta Bobola, Neil A. Hanley

## Abstract

How the genome activates or silences transcriptional programmes governs organ formation. Little is known in human embryos undermining our ability to benchmark the fidelity of in vitro stem cell differentiation or cell programming, or interpret the pathogenicity of noncoding variation. Here, we studied histone modifications across thirteen tissues during human organogenesis. We integrated the data with transcription to build the first overview of how the human genome differentially regulates alternative organ fates including by repression. Promoters from nearly 20,000 genes partitioned into discrete states without showing bivalency. Key developmental gene sets were actively repressed outside of the appropriate organ. Candidate enhancers, functional in zebrafish, allowed imputation of tissue-specific and shared patterns of transcription factor binding. Overlaying more than 700 noncoding mutations from patients with developmental disorders allowed correlation to unanticipated target genes. Taken together, the data provide a new, comprehensive genomic framework for investigating normal and abnormal human development.

## INTRODUCTION

Organogenesis is the key phase when the body’s tissues and organs are first assembled from rudimentary progenitor cells. In human embryos this is the critical period during weeks five to eight of gestation when disruption can lead to major developmental disorders. While approximately 35% of developmental disorders are explained by damaging genetic variation within the exons of protein-coding genes^1^, de novo mutation (DNM) in the noncoding genome has been associated increasingly with major developmental disorders^2^. The noncoding genome also harbours over 80% of single nucleotide polymorphisms (SNPs) implicated in genome wide association studies (GWAS) for developmental disorders, or in GWAS of later onset disease, such as schizophrenia and type 2 diabetes, where contribution is predicted from early development^3^. These genetic alterations are presumed to lie in enhancers for developmental genes or in other regulatory elements such as promoters for noncoding RNAs that may only be active in the relevant tissue at the appropriate stage of organogenesis. Aside from rare examples^4^, this has remained unproven because of lack of data in human embryos. While regulatory data are available in other species at comparable stages^5^, extrapolation is of limited value because the precise genomic locations of enhancers are poorly conserved^6,7^ even allowing for enriched sequence conservation around developmental genes^8,9^. Sequence conservation alone is also uninformative for when and in what tissue a putative enhancer might function. Comprehensive regulatory information is available from later fetal development via initiatives such as NIH Roadmap^10^ but these later stages largely reflect terminally differentiated, albeit immature cells rather than progenitors responsible for organ formation. In contrast, a small number of studies on a handful of isolated tissues, such as limb bud^11^, craniofacial processes^12^, pancreas^13^ or brain^14^, have demarcated regulatory elements directly during human organogenesis. However, most organs remain unexplored. Moreover, nothing is known about patterns of regulation deployed across tissues, which is an important factor because tissues are often co-affected in developmental disorders. To address these gaps in our knowledge we set out to build maps of genome regulation integrated with transcription during human organogenesis at comprehensiveness currently unattainable from single cell analysis.

## RESULTS

Organs and tissues from thirteen sites were microdissected and subjected to chromatin immunoprecipitation followed by deep sequencing (ChIPseq) for three histone modifications (Figure 1a): H3K4me3, enriched at promoters of transcribed genes; H3K27ac, at active enhancers and some promoters; and H3K27me3 delineating regions of the genome under active repression by Polycomb. Tiny tissue size and the scarcity of human embryonic tissue required some pooling and precluded study of additional modifications. Biological replicates were undertaken for all but two tissue sites (Supplementary Table 1). Tissues and stages were matched to polyadenylated RNAseq datasets acquired at sufficient read depth to identify over 6,000 loci with previously unannotated transcription (Supplementary table 2)^15^. Overlaying the data revealed characteristic tissue-specific patterns of promoter and putative enhancer activity, and novel human embryonic transcripts. This was particularly noticeable surrounding genes encoding key developmental transcription factors (TFs), such as the example shown for *NKX2-5* in heart (Figure 1b). Tissues lacking expression of the TF gene tended to carry active H3K27me3 modification (rather than simply lack marks). Putative tissue-specific enhancer marks were characteristically distributed over several hundred kilobases (heart-specific example to the far right of Figure 1b). These isolated H3K27ac marks were often unpredicted by publically available data from cell lines or terminally differentiated lineages and were not necessarily conserved across vertebrates (mean per-base phyloP score 0.175; range - 1.42 to +6.94 for n=51,559 regions)^16^. Unexpected H3K4me3 and H3K27ac peaks that failed to map to the transcriptional start sites (TSSs) of annotated genes mapped to the TSS of novel human embryo-enriched transcripts, such as the bidirectional *HE-TUCP-C5T408* and *HE-LINC-C5T409* (Figure 1b; see Supplementary File 1H in reference^15^ for the complete catalogue).

**Figure 1.**
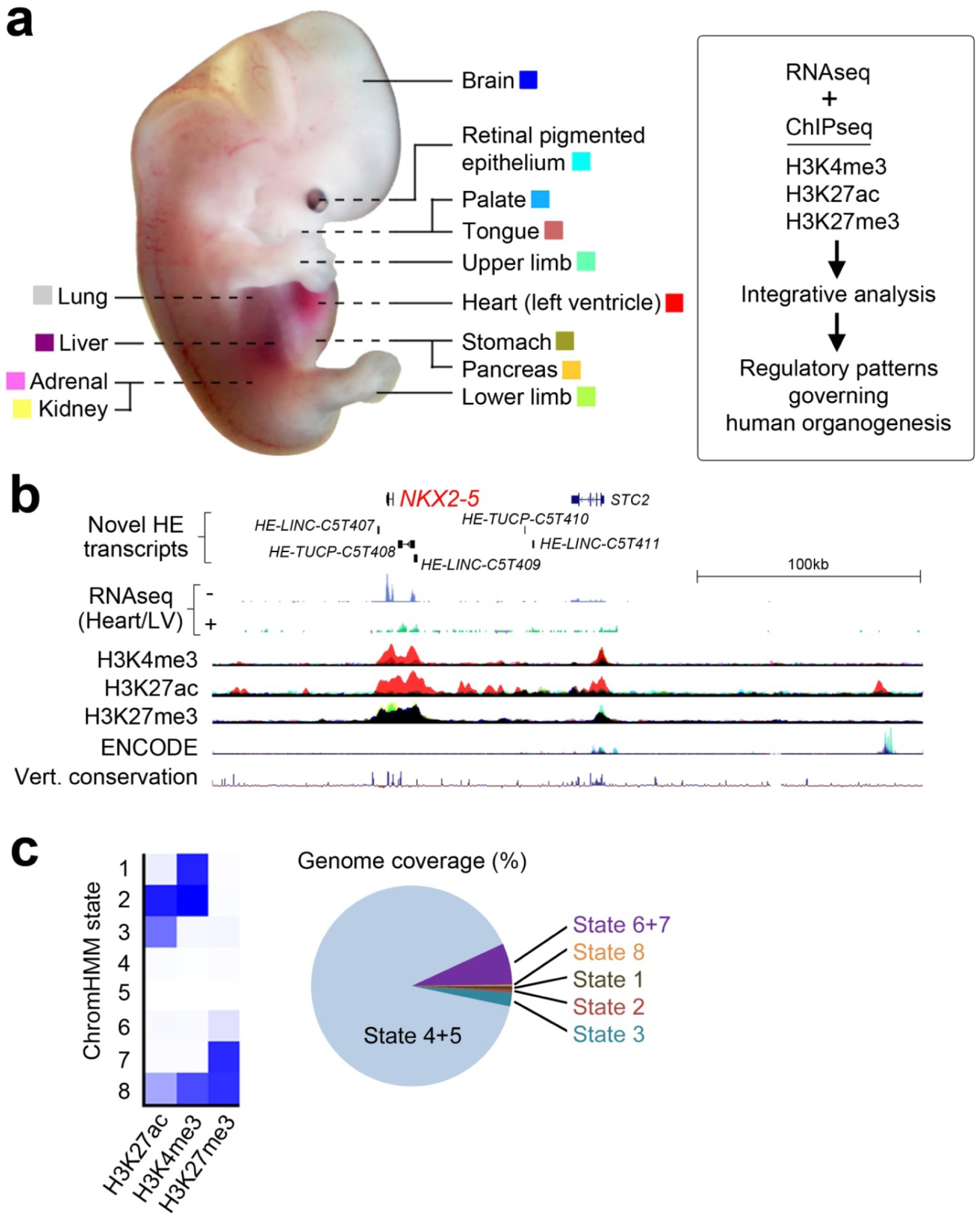
Epigenomic landscape across thirteen human embryonic tissues. a) Thirteen different human embryonic sites were sampled for RNAseq^15^ and ChIPseq as described in the Materials and Methods and in Supplementary tables 1 and 2. The same colour coding for each tissue is applied throughout the manuscript in overlaid ChIPseq tracks. The heart (left ventricle) dataset is summarised as ‘Heart/LV’ from hereon. b) 300 kb locus around the *NKX2-5* gene, the most discriminatory TF gene for human embryonic heart^15^. The locus contains five novel human embryonic (*HE*) transcripts enriched in heart [three *LINC* RNAs and two transcripts of uncertain coding potential (*TUCP*)]. Heart/LV-specific (red) H3K4me3 and H3K27ac marks were detected at the *NKX2-5* TSS and adjacent novel transcripts (*HE-TUCP-C5T408* and *HE-LINC-C5T409*). Novel heart-specific H3K27ac marks were visible up to 200 kb away (e.g. at the extreme right of panel). H3K27me3 marked the region from *NKX2-5* to *HE-LINC-C5T409* in all non-heart tissues (the track appears black from the superimposition of all the different colours other than red). ENCODE data are from seven cell lines^23^. c) Genome coverage by ChromHMM for the different histone modifications was similar across all tissues (Supplementary figure 1) with an average 89.8% of the genome unmarked (range: 81.7-94.0; States 4 & 5) and 3.3% consistent with being an active promoter and/or enhancer (range: 1.7-6.1; States 1-3).

By analysis based on a Hidden Markov Model the genome partitioned into different chromatin states very similarly across tissues^17^. While three histone marks allowed for eight different segmentations, aggregation into fewer states was possible (Figure 1c). On average across tissues, 3.3% of the genome was active promoter (States 1 & 2; H3K4me3 +/- H3K27ac) or putative enhancer (State 3; H3K27ac) (range 1.7-6.1%; Figure 1c & Supplementary figure 1). 6.7% was variably marked as actively repressed (States 6 & 7; range 3.3-13.0; H3K27me3), while on average 89.8% of the genome was effectively unmarked (States 4 & 5; range 81.7-94.0). ∼0.2% seemingly had both H3K4me3 and H3K27me3 marks with some detection of H3K27ac (State 8; range 0.16-0.33). This latter state has been considered bivalent and characteristic of ‘poised’ genes whose imminent expression then initiates cell differentiation pathways^18-20^. Ascribing bivalency has been reliant on setting an arbitrary threshold for whether a site is marked or not, and might simply reflect mixed marks due to heterogeneity in a cell population. To avoid the need for thresholding we clustered promoter profiles for each histone mark integrated with transcription over 3 kb either side of 19,791 distinct protein-coding TSS in each tissue^21^ (Figure 2 and Supplementary figures 2-4). Broader H3K4me3 and H3K27ac signals at the TSS correlated with higher levels of transcription (Figure 2a-b; we termed this promoter state ‘Broad’ expressed versus ‘Narrow’ or ‘Bi-directional’ expressed). 25-30% of genes across tissues were unmarked and lacked appreciable transcription (‘Inactive’). These promoters typically lacked CpG islands (<20% compared to 67.7% of the 19,791 genes). Conversely, 90-95% of TSS regions marked with H3K27me3 featured CpG islands with an over-representation of TFs; 31.2% of TFs (n=1,659) were actively repressed in at least one tissue compared to 20.0% of non-TF genes (odds ratio 1.82, confidence interval 1.63-2.04; p-value <2.2e-16). H3K27me3 detection at the TSS was ∼50% greater for genes encoding TFs (Supplementary figure 5). H3K27me3 was only detected with minimal accompaniment of H3K4me3 or H3K27ac and no transcription (Figure 2 a-b). The categorization for each of the 19,791 genes in all tissues is listed in Supplementary table 3. This neat partitioning would not have been possible if the data were overly confounded by cellular heterogeneity. The data argue against a major bivalent chromatin state at gene promoters in progenitor cells during human organogenesis.

**Figure 2.**
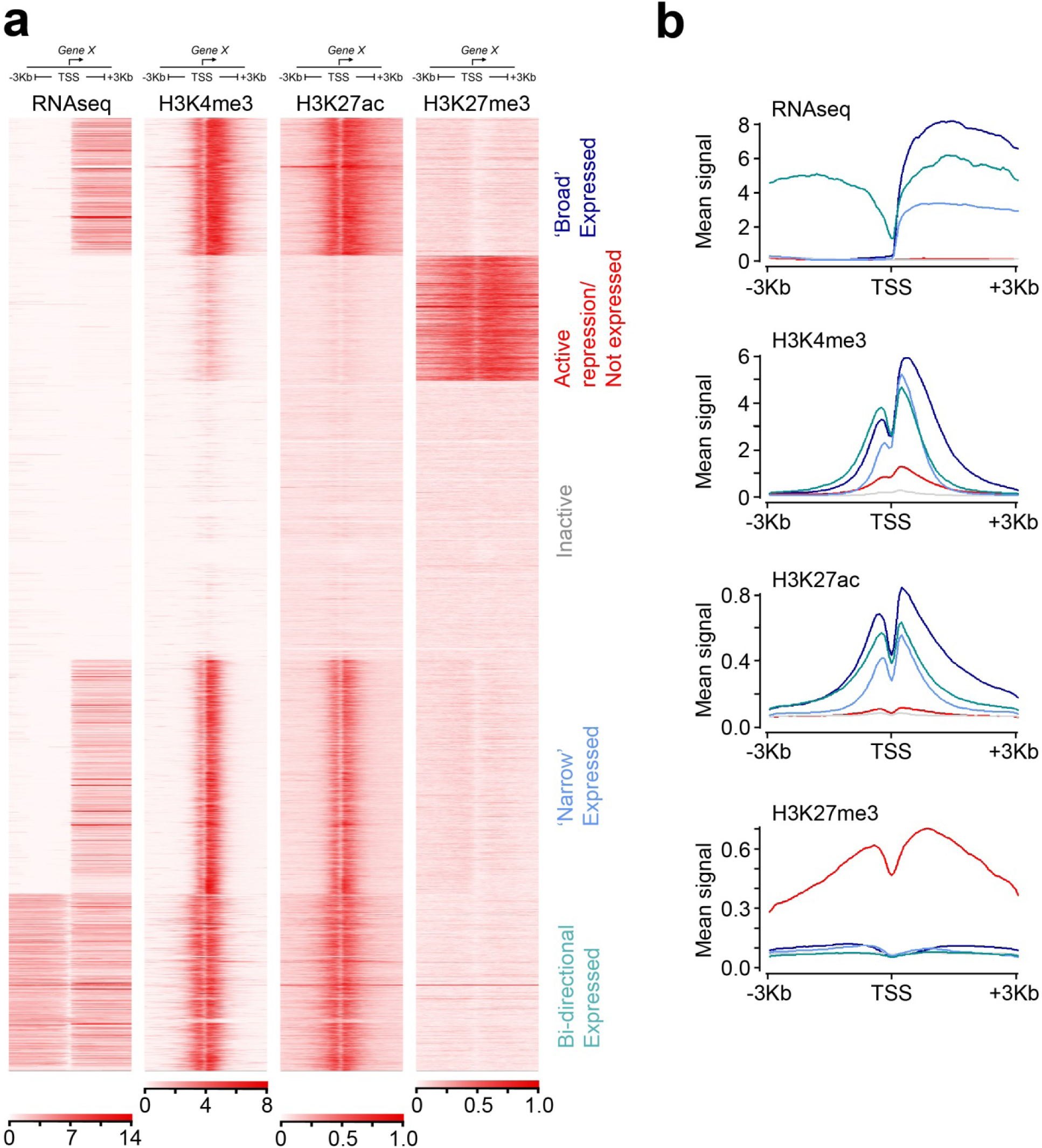
Classification of genes into discrete states according to characteristics at the promoter associated with transcription. a) Clustered heatmaps surrounding the transcriptional start sites (TSS +/- 3 kb) of 19,791 annotated genes. The example shown is for adrenal. One replicate is shown for each data-type for simplicity. Replicates across all tissues were near identical. Two minor variations on this pattern were detected in RPE (Supplementary figure 2) and liver, lung and brain (Supplementary figure 3). b) Mean signal levels for the genes clustered in a). Traces are coloured according to the text colour in a). ‘Broad expressed’ genes show approximately double the level of transcription and twice the width of H3K4me3 and H3K27ac marks compared to ‘Narrow expressed’ genes.

This classification allowed us to ask how promoter state changed across different tissues. Tracking all states in all tissues was complex to visualise (Supplementary figure 6). Unifying ‘Broad’, ‘Narrow’ and ‘Bi-directional’ expressed into a single category (‘Expressed’) displayed how the majority of genes remained unaltered across the thirteen tissues (Figure 3a). In contrast, 29% of genes had a variable promoter state. Within this subset we predicted that genes responsible for a specific organ’s assembly, such as developmental TFs, would need to be actively excluded or ‘disallowed’ at inappropriate sites (as seen in Figure 1b for *NKX2-5*). We tested this in the replicated datasets by comparing genes transcribed uniquely in one tissue for either inactivity (no mark) or active repression elsewhere (H3K27me3; disallowed). Gene ontology (GO) analysis of the ‘uniquely expressed/disallowed elsewhere’ gene sets identified the appropriate developmental programme in all instances (as shown for heart in Figure 3b; e.g. ‘heart development’). In contrast, tissue-specific transcription initiated from genes that were simply inactive in other organs tended to highlight differentiated cell function (Figure 3b; e.g. sarcomere organization). These observations highlight the preferential use of H3K27me3 at the promoters of genes controlling cell fate decisions but not differentiated function. To study this over time, we included datasets from human pluripotent stem cells (hPSCs) and adult tissue, to scrutinize regulatory changes temporally. Different sets of repressed genes lost their H3K27me3 mark to become expressed as cells transitioned from pluripotency to embryonic pancreatic progenitors or from pancreatic progenitors to mature pancreas (Figure 3c). Surprisingly, the same KEGG term relating to monogenic diabetes emerged in both instances (Figure 3d). However, the genes underlying the first transition related to early function in pancreatic organogenesis, hypoplasia or aplasia; while the genes in the second transition specifically related to post-embryonic pancreatic islet cell differentiation and beta-cell function^22^.

**Figure 3.**
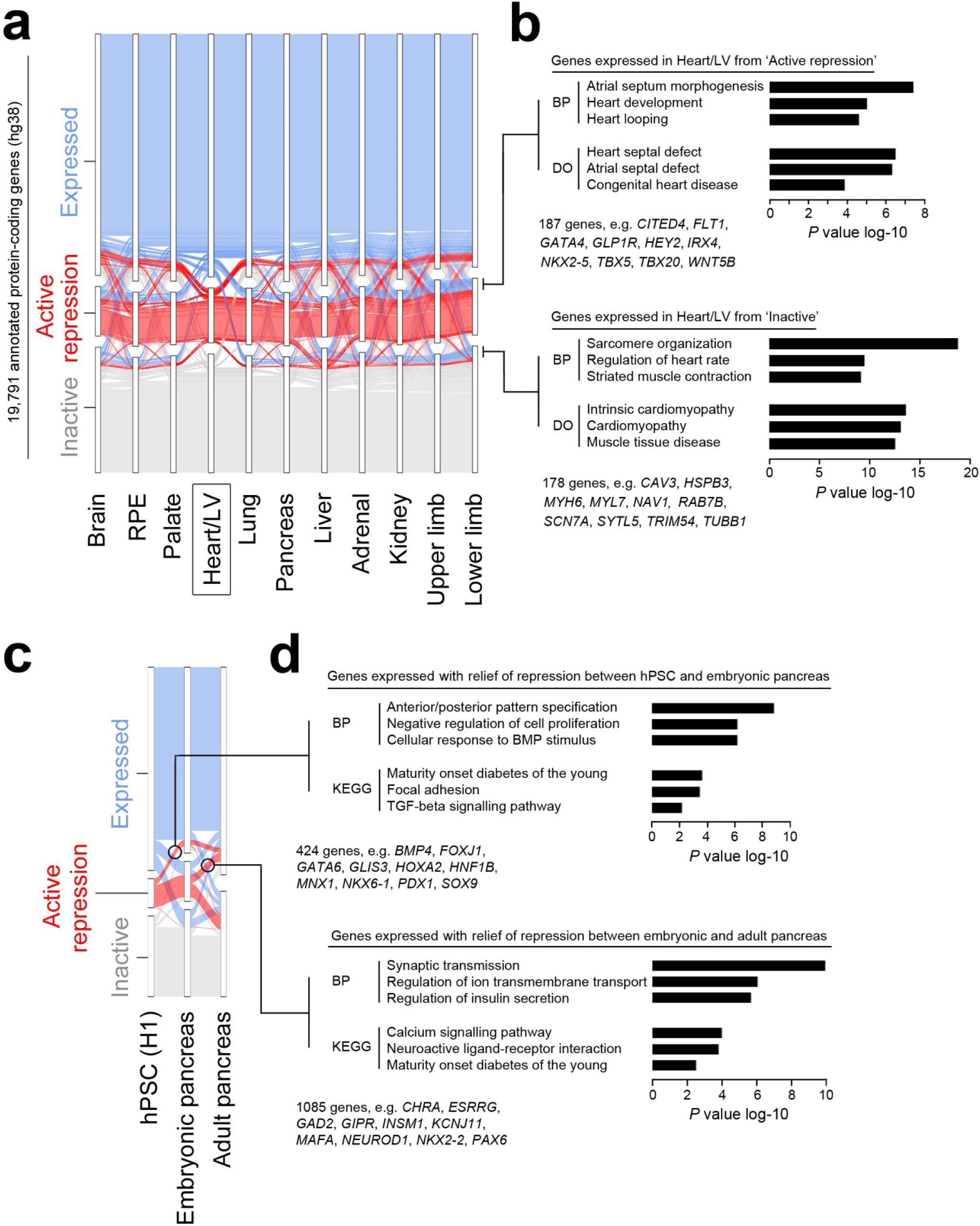
Integration of promoter states across tissues and over time to decipher developmental programmes and associated disease in individual organs. a) Alluvial plot showing promoter state for 19,791 annotated genes across all tissues with replicated datasets. To aid visualisation all the different transcribed states are amalgamated into a single ‘Expressed’ category (the alluvial plot for all individual states is shown in Supplementary figure 6). The example shown is centred on the promoter state in the Heart/LV dataset. Those genes with an ‘Expressed’ promoter state in heart and either ‘Active repression’ or ‘Inactive’ elsewhere are indicated to the right of the panel and subject to gene enrichment analyses in b). b) Gene enrichment analysis of genes with an ‘Expressed’ promoter state in heart and either ‘Active repression’ or ‘Inactive’ in all remaining tissues. Examples of the genes underlying the biological process (BP) or disease ontology (DO) terms and their total number are listed beneath the bar charts. c) Alluvial plot showing the variance in promoter state between H1 human pluripotent stem cells (hPSCs), the embryonic pancreas (prior to endocrine differentiation^22^) and the adult pancreas. Circles capture those genes that shift from ‘Active repression’ to ‘Expressed’ at the stage of either embryonic or adult pancreas. d) Gene enrichment analyses of encircled genes from c). Examples of the genes underlying the BP and KEGG terms and their total number are listed beneath the bar charts. While maturity onset diabetes of the young emerged in both analyses, the underlying genes were different reflecting developmental roles prior to or after pancreatic endocrine differentiation^22^.

Having recognized the disallowed status of developmental TFs in inappropriate tissues, we wanted to test whether our putative intergenic human embryo-enriched enhancers were capable of driving appropriate reporter gene expression at the correct locations in developing zebrafish. We identified H3K27ac marks that were enriched in the human embryo compared to 161 ENCODE or NIH Roadmap datasets^10,23^ and not detected in the FANTOM5 project^24^. We developed an algorithm to test for embryonic tissue specificity and filtered for sequence conservation (not necessarily in zebrafish; see Materials & Methods). We manually inspected the remainder for proximity (<1 mb) to genes encoding TFs and, in particular, to increase clinical relevance, to those associated with major developmental disorders. We ensured no H3K4me3 or polyadenylated transcription in the immediate vicinity (i.e. an unannotated promoter). We tested ten such enhancers out of 44 within 1 mb of *TBX15, HEY2, ALX1, IRX4, PITX2, HOXD13, NKX2-5, WT1, SOX11* and *SOX9* for their ability to direct appropriate GFP expression in stable lines of transgenic zebrafish (Supplementary table 4). Two (h-003-kid near *WT1* and h-022-mix near *SOX11*) failed to generate any GFP in any location. The remaining eight all yielded GFP at the predicted site in zebrafish embryos (Figure 4 and Supplementary table 4), despite only one of the putative enhancer sequences being conserved in zebrafish (Fig. 4a, h-027-lim near *TBX15*). These data imply that our H3K27ac detection marks novel functional enhancers operating over considerable distance. Moreover, inter-species sequence conservation is not needed for appropriate reporter gene expression (i.e. it is the TFs which bind that are conserved).

**Figure 4.**
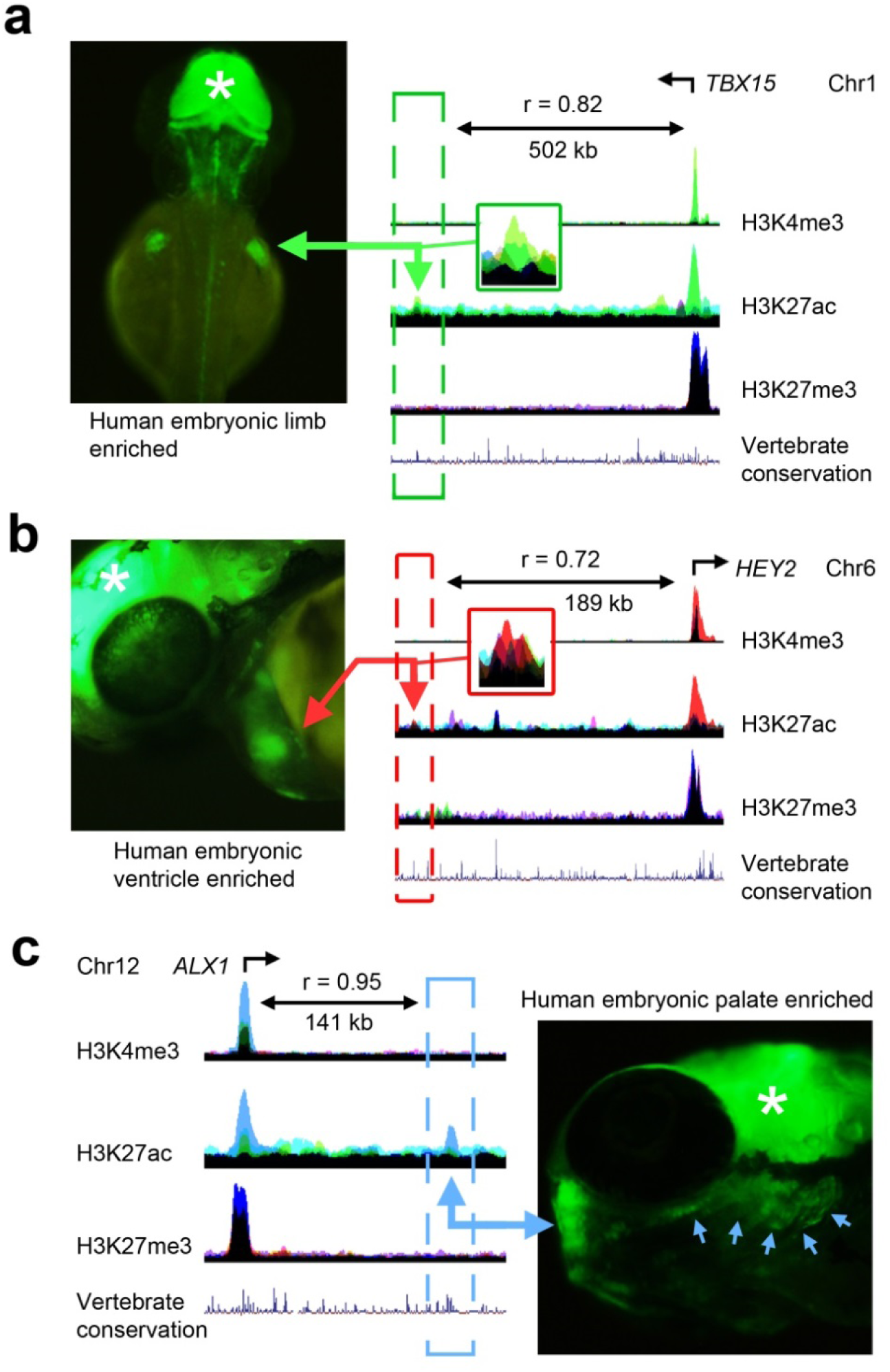
Stable transgenic analysis in developing zebrafish of regions marked by H3K27ac in human embryonic tissues. H3K27ac marked regions were tested in multiple lines of stable transgenic zebrafish (details in Supplementary table 4; same colour coding of tracks as in Figure 1). a) 231 bp limb enhancer, 502 kb downstream of *TBX15*, with corresponding green fluorescent protein (GFP) detection in fin bud at 48 hours post-fertilisation (hpf). b) 355 bp heart/LV enhancer, 189 kb upstream of *HEY2*, with corresponding ventricular GFP detection at 48 hpf. c) 1.5 kb palate enhancer, 141 kb downstream of *ALX1*, with GFP in the developing trabecula and mandible (blue arrows) at 48 hpf. Correlations between the enhancer and transcription of the transcription factor are shown for each example. Note the H3K27me3 marks over the gene in each instance in other tissues. *, midbrain GFP expression from the integral enhancer in the reporter vector used as positive control for transgenesis.

We wanted to explore the link between these regulatory elements and surrounding gene expression at genome-wide scale. Assured that ChIPseq marks were reproducible within biological replicates without batch effect (Supplementary figure 7) we parsed the genome into 3,087,584 non-overlapping 1 kb bins. Reads within each bin were counted for each mark. Phi correlation between biological replicates indicated this approach to peak calling was very similar to using MACS (Supplementary figure 8). Counts were downsampled and averaged within tissues and correlated with RNAseq data from the same tissue over 1 mb in either direction (i.e. a 2 mb window). On average this window included 44 annotated genes (range: 0-247 genes). For those H3K27ac marks which functioned in zebrafish the strongest correlation was with the appropriate TF gene, for instance *TBX15* over ∼500 kb in limb (Figure 4a). Moreover, different H3K27ac marks could be correlated to the same gene, potentially allowing previously unknown enhancers to be grouped, for instance in the adrenal around the adrenal hypoplasia gene, *NR0B1*, located on the X chromosome (Supplementary figure 9).

Parsing the ChIPseq data into bins allowed integration of information across tissues, which is challenging when based on empirical modelling by MACS. Placing raw read counts per bin in rank order produced near identical ‘elbow’ plots for all marks in all tissues. This allowed the point of maximum flexure to be used quantitatively for calling marks in a binary ‘yes/no’ fashion (Figure 5a). This simplified calling facilitated exploration of regulatory patterns across tissues. Requiring a bin to be marked in any two or more samples identified 48,570 different H3K27ac patterns genome-wide. By genome coverage all tissue-specific patterns ranked within the top one percent (Figure 5b; heart came first). H3K27ac showed far more tissue selective patterns than H3K4me3 or H3K27me3 (Figure 4b and Supplementary figures 10-11). Motif analysis on the tissue-specific H3K27ac regions allowed imputation of master TFs for individual tissues, such as NR5A1 in 54.5% of adrenal-specific bins (n=18,411) compared to 25% of the remaining 141,706 bins (Figure 5c). Mutation of *NR5A1* causes adrenal agenesis in human and mouse (OMIM 184757). MEF, TBX and bHLH family members emerged in the heart-specific bins (Figure 5c); all are associated with congenital heart disease^25^. Having integrated our data we could also uncover regulatory regions that were shared precisely across two or more tissues to explore developmental disorders which manifest in multiple organs. Novel enrichment for composite PITX1/bHLH motifs was found in limb and palate (Figure 5c). GATA binding motifs were enriched in heart and pancreas. Shared patterns could be explicitly instructed by requiring detection in four or more samples (Supplementary figure 12). We hypothesized that patterns shared across many tissues ought to contain elements regulating generic developmental functions. Scrutinising bins marked in over half of all H3K27ac samples (n=30,226 bins versus remaining background of 80,352 bins) identified enrichment for the ETS motif. ETS transcription factors are involved in cell cycle control and proliferation^26^.

**Figure 5.**
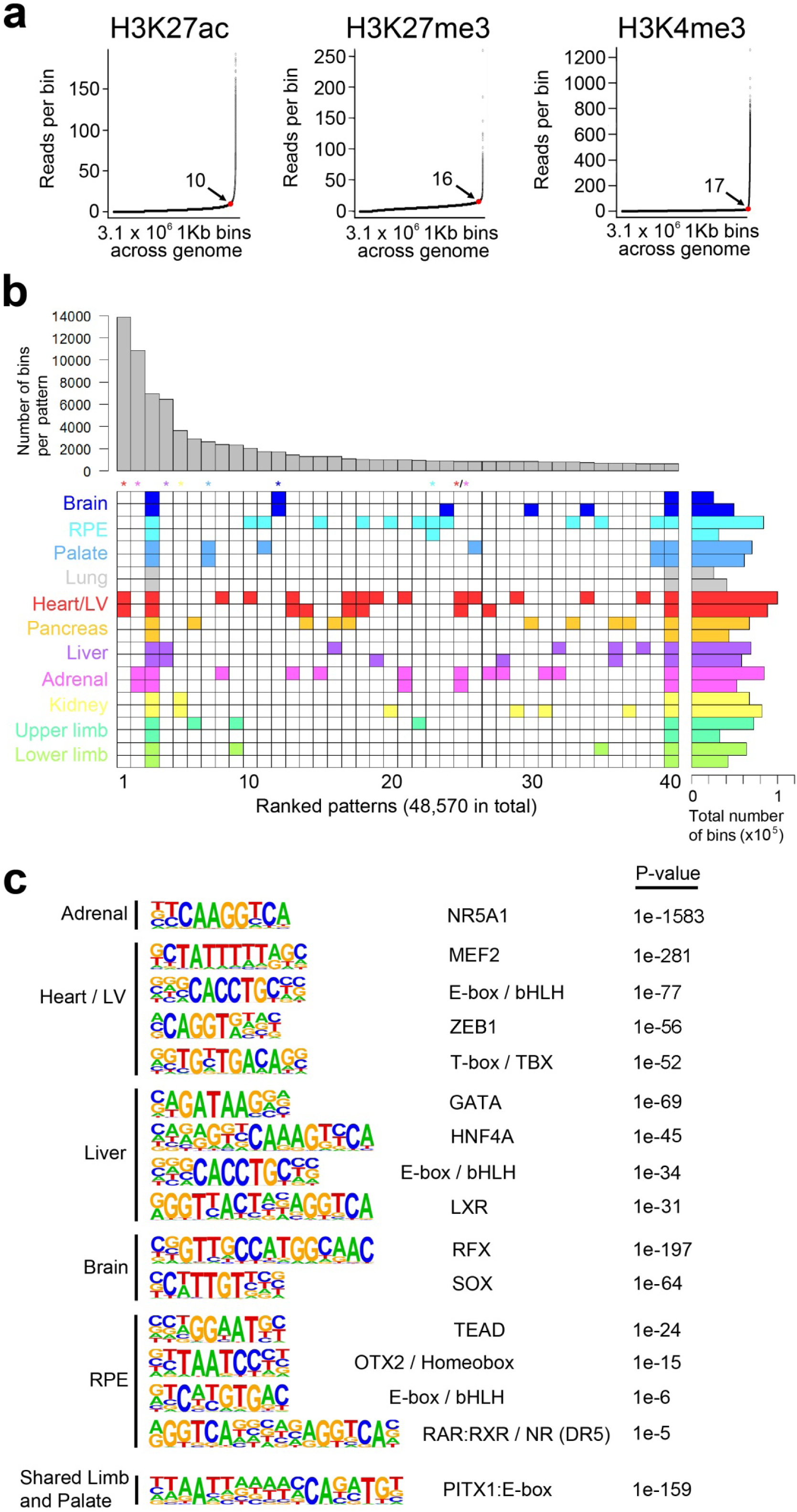
Imputing patterns of enhancer activity and regulation by developmental transcription factors across human embryonic tissues. a) Elbow plots for each histone modification following allocation of the genome into 3.1 million consecutive bins of 1 kb. The example shown is for adrenal providing the number of reads per bin at the point of maximum gradient change (the ‘elbow point’, red dot) and a quantitative measure of whether a bin was marked or not (e.g. >10 or <10 respectively for H3K27ac). Converting marks into a binary ‘yes/no’ call at any point in the genome facilitated data integration across the different tissues. While the number of reads per bin at the elbow point was different for each mark across the tissues the shape of the curve remained the same. b) Euler grid for bins marked by H3K27ac (defined by elbow plots) in replicated tissues (i.e. two rows/replicates per tissue). Total number of marked bins per individual dataset is shown to the right. The example in b) required a bin to be called in any two or more samples and is ordered by decreasing bin count per pattern (bar chart above the grid). A total of 48,570 different patterns were identified of which the top 40 are shown. Tissue-specificity for all sites emerged in the top 265 (0.5%) patterns; colour-coded asterisks above columns). For example, nearly 14,000 bins marked only in the two Heart/LV H3K27ac datasets ranked first as the most frequent pattern. The seventh most frequent pattern in nearly 3,000 bins was palate-specific. Tissue-specific patterns were far less apparent at promoters (H3K4me3, n=18,432; Supplementary figure 10) or for H3K27me3 (n=26,339; Supplementary figure 11). While patterns across multiple tissues were permitted by stipulating marks in ≥ 2 samples (e.g. heart and adrenal in column 24), they could be enforced by stipulating marks in at least four samples (Supplementary figure 12). c) Enrichment of known TF-binding motifs in the tissue-specific patterns of H3K27ac identified in b). Five individual tissues are shown as examples alongside analysis of the shared regulatory pattern identified for limb and palate identifying marked enrichment of a compound PITX1:E-box motif.

Noncoding mutations in promoters or enhancers have been linked increasingly to major developmental disorders^4,27^. Previously, as part of the Deciphering Developmental Disorders (DDD) study, we studied 7,930 individuals and their parents^28^. 87% of patients had neurodevelopmental disorders. 10% had congenital heart defects. 68% of patients lacked disease-associated DNMs within exomes (‘exome-negative’) pointing to the likely importance of the noncoding genome^2^. We sequenced 6,139 non-coding regions (4.2 Mb) selected as ultra-conserved regions (UCRs: n=4,307), experimentally validated enhancers (EVEs: n=595) or as putative heart enhancers (PHE: n=1,237) and found 739 non-coding DNMs^2^. 78% of the 6,139 regions were marked by H3K27ac or H3K4me3 in our embryonic tissues, with a higher percentage overlap for the EVEs (87%) and near perfect overlap for the PHEs (99%) (Figure 6a). An additional 9% were marked by H3K27me3 suggesting non-coding regulation in a currently unsampled tissue. The distribution of DNMs was very similar (Figure 6b). Nearly half of the regions containing DNMs were marked by H3K27ac and/or H3K4me3 that was replicated in at least one tissue. Most commonly, this included the heart or brain, in keeping with the predominance of neurodevelopmental and cardiac phenotypes in the DDD cohort and the PHEs selected for sequencing (Figure 6c). 75% of the PHEs with DNMs mapped to replicated H3K27ac and/or H3K4me3 in our heart dataset. This rose to 100% if the need for replication was removed. We did not observe enrichment for DNMs in patients with heart or limb phenotypes in elements marked by H3K27ac or H3K4me3; but the power of this test was most likely hampered by low patient numbers (Figure 6d). Enrichment for DNMs in elements marked by H3K27ac was detected for the greater number of cases with neurodevelopmental disorders (1.45-fold, 95% confidence interval 1.09-1.90; p=0.0056) (Figure 6d). This was similar to our previous report using NIH Roadmap H3K27ac and/or DNaseI hypersensitivity data derived from second trimester fetal brain^2^. Our results support a role for non-coding mutations in severe neurodevelopmental disorders and that regulatory marks active during human organogenesis could help stratify disease-relevant non-coding regions.

**Figure 6.**
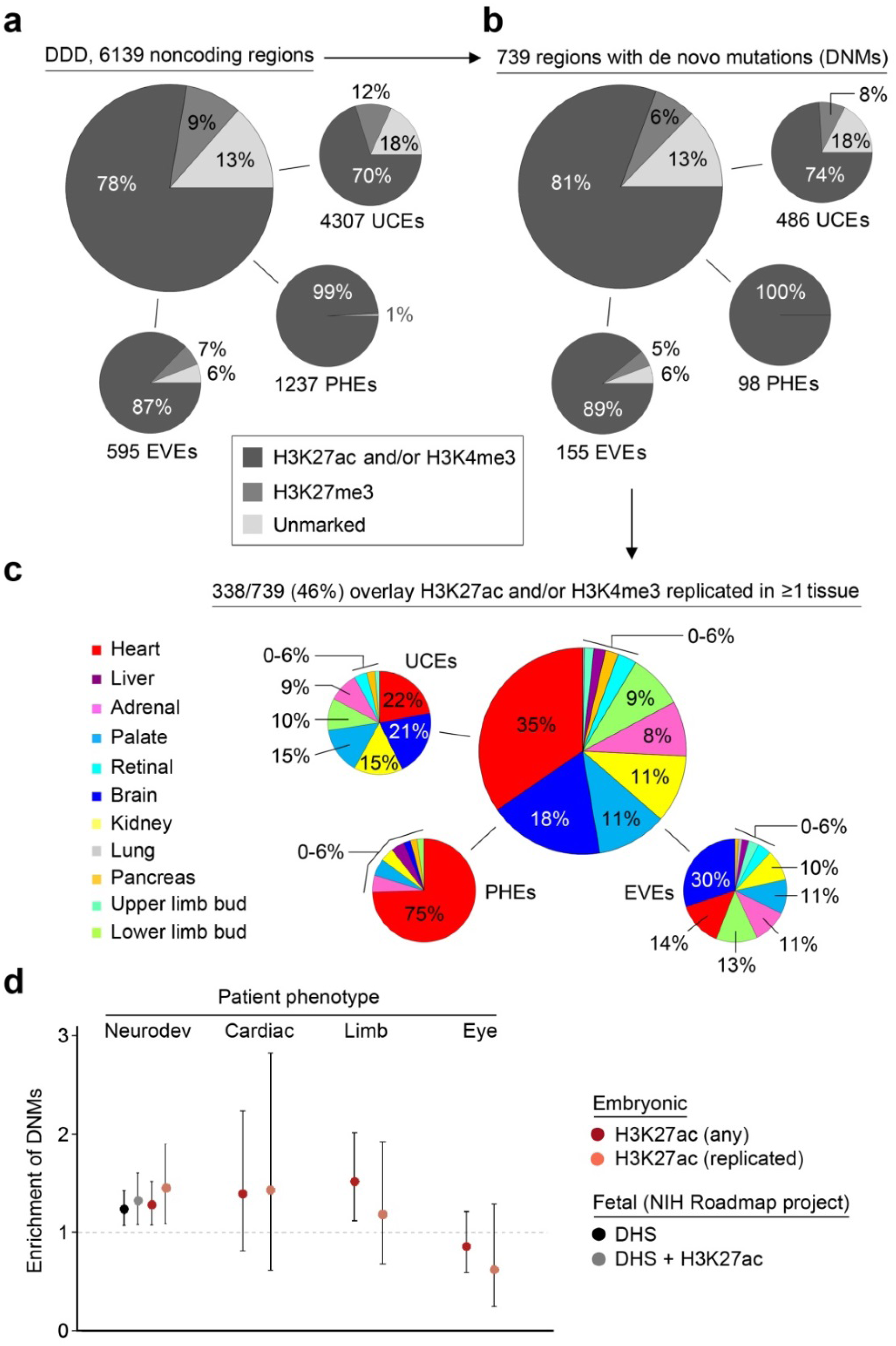
Intersection of the epigenomic landscape during human organogenesis with noncoding de novo mutations linked to developmental disorders in patients. a) The Deciphering Developmental Disorders (DDD) study included 6,139 noncoding regions in its sequence analysis of trios comprising affected individuals and unaffected parents^2,28^. These noncoding regions were selected on the basis of high sequence conservation (ultra-conserved elements, UCEs, n=4,307), experimental validation (experimentally validated enhancers, EVEs, n=595) or identification as a putative heart enhancer (PHE, n=1,237). Overlap with any H3K27ac, H3K4me3 or H3K27me3 1 kb bins is shown as an aggregate and for each individual category (UCE, EVE or PHE). b) Equivalent overlap is shown for the 739 regions in which disease-associated de novo mutations (DNMs) were identified. c) 46% of DNM-positive regions were situated (+/- 1 kb) in at least one tissue-replicated H3K27ac and / or H3K4me3 bin. Over half of the disease-associated overlap was covered by heart/LV (35%) and brain (18%). 75% of the disease-associated PHE regions were situated within 1kb of a heart/LV-specific histone mark. d) Enrichment (observed/expected) in the number of DNMs overlapping (+/- 1 kb) H3K27ac marks during human organogenesis. For neurodevelopmental phenotypes this included analysis against DNAse hypersensitivity data and H3K27ac data from second trimester fetal brain^10^. Error bars show the 95% confidence limits for the mean calculated for a Poisson distribution^52^.

Prioritising DNMs for potential pathogenicity and how they might disrupt surrounding gene function is very challenging. For developmental disorders our mapping allowed focus on enhancers and promoters in the relevant tissue at an appropriate embryonic stage. Our comprehensive tissue-by-tissue catalogue of transcription also allowed more detailed consideration of DNMs in close proximity to previously unappreciated human embryonic noncoding RNAs. Correlating histone modifications with gene expression across all tissues offered a means of prioritising target gene(s). As an example, we identified a G-to-T DNM in a patient with a neurodevelopmental disorder in a UCR on chromosome 16 (Figure 7; chr16:72,427,838). The mutation is situated in the middle of the annotated LINC RNA, *LINC01572*, expressed in testis. Our data illustrated numerous surrounding human embryonic noncoding transcripts. In fact, the DNM was located at the TSS of *HE-OT-AC004158.3*, expressed at 19.5-fold higher levels in human embryonic brain than any other tissue (mean read count of quantile normalized transcripts in brain, 1317.2; mean in other tissues, 32.2), in a 4 kb region of brain-specific H3K27ac (and to a lesser extent, H3K4me3). Amongst at least 18 protein-coding genes in the surrounding region, the H3K27ac signal was most highly correlated to expression of *ZNF821* (r=0.92) located approximately 550 kb away and anticorrelated to expression of the adjacent gene, *ATXN1L* (r=-0.65; Figure 7). Taken together, these data and correlations, available to browse as tracks on the UCSC Genome Browser, build a human embryonic atlas of developmental regulatory information linked to gene expression for the overlay of variants identified by clinical sequencing and GWAS.

**Figure 7.**
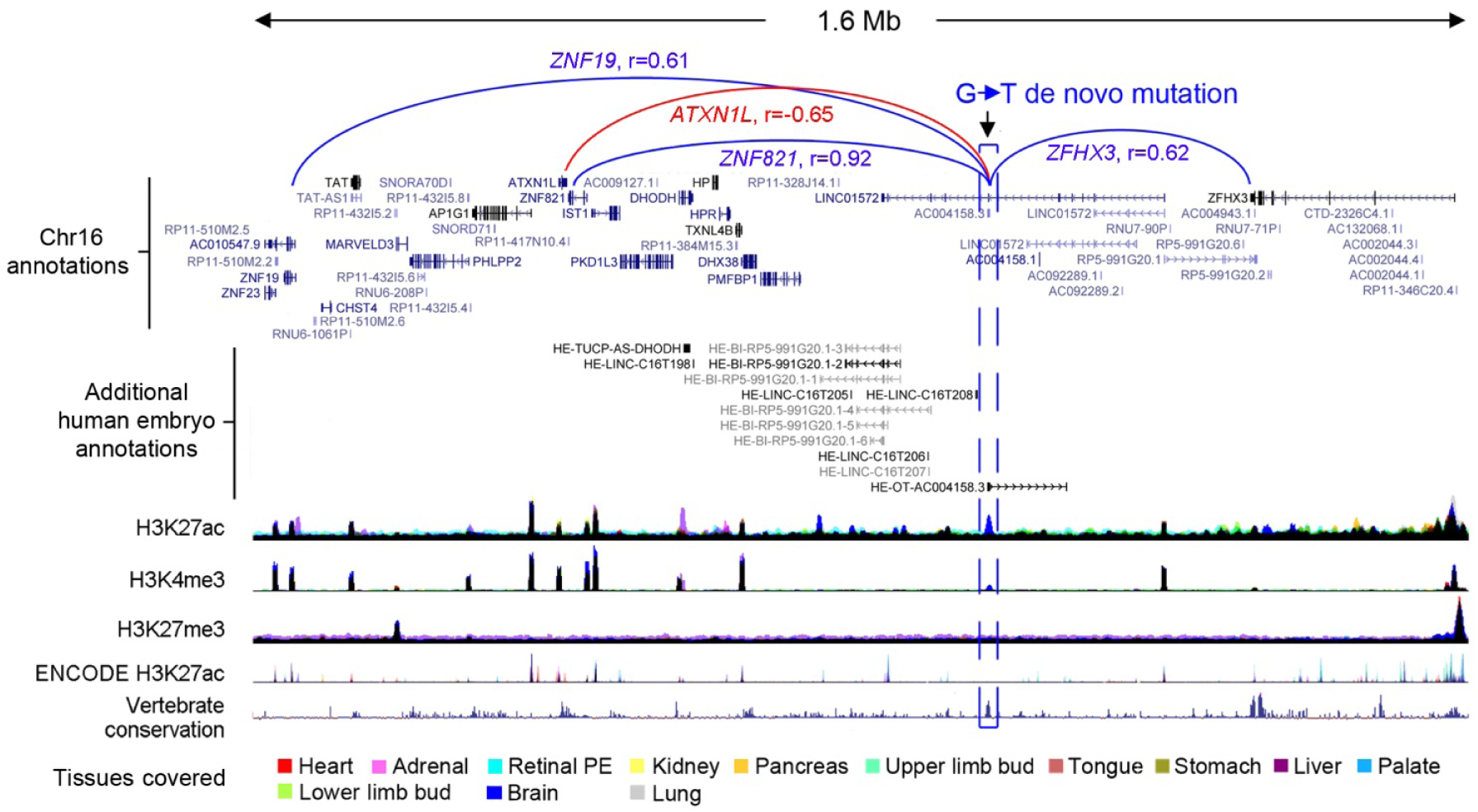
Overlap of individual disease-associated de novo mutations with the human embryonic epigenome correlated to surrounding gene expression. An intergenic G-to-T de novo mutation (DNM; hg38, chr16:72427838) is shown for a patient with a neurodevelopmental phenotype. Tracks are shown demonstrating novel human embryonic noncoding transcription (enriched in human embryonic brain), the three epigenomic marks, ENCODE data^23^ and conservation amongst vertebrates. The DNM overlaps a brain-specific (dark blue) H3K27ac and small H3K4me3 mark. The highest correlations are shown, notably to the promotors of *ZNF821* (r=0.92) (dark blue) with anticorrelation (r=-0.65, red) to the adjacent gene, *ATXN1L.*

## DISCUSSION

Previous studies of enhancer usage in human embryos have tended to focus on individual tissues inferring, amongst other findings, aspects of genome regulation responsible for human-specific attributes^11-14^. Here, we incorporated epigenomic data with transcription across thirteen sites during human organogenesis to build tissue-by-tissue maps of enhancers and promoters linked to gene expression. While similar to prior work in mouse^5^ and building on our previous transcriptomic atlas^15^, the integrated approach here offers new opportunities to understand how human organ formation is regulated in health and disease.

As cost has declined, whole genome sequencing (WGS) has become an important tool in main stream clinical investigation, opening up potential genetic diagnoses in the 98.5% of the human genome that lies outside of coding sequences. However, assessing the non-coding genome is very challenging: millions of rare variants are returned in each individual, while only one might be pathogenic^29^. Functional analysis, even of a handful of variants, is clinically impractical. For non-coding mutations to affect organogenesis (either in developmental disorders or in later life disease such as type 2 diabetes where there is an embryonic contribution) it is logical that mutations are located in regulatory regions of the genome that are active in post-implantation human embryos. As evident from Figure 1c, our identification of this landscape offers a timely, new pipeline for stratifying 98.5% of the genome down to 3% on average per tissue (States 1-3). Enrichment of tissue-specific TF binding in these enhancers and promoters reinforced our previous findings based solely on computational analysis of 5’ flanking regions for the importance of NR5A1 in adrenal and HNF4A in liver^15^. However, the integrated sampling of numerous sites uncovered far more complex patterns of regulation operating across tissues. The enrichment of *PITX1* binding motifs in active regulatory regions uniquely shared across limb bud and palate fits with mutations in *PITX1* causing limb defects and cleft palate^30^. Similarly, GATA4 and GATA6, inferred from regulatory regions shared uniquely between heart and pancreas, are the only two TFs linked to the dual phenotype of cardiac malformation and monogenic diabetes^31,32^. Overlaying GWAS data with chromosomal conformation studies from older human fetal brain has prioritized target genes for risk of schizophrenia^33^. While these techniques are yet to be applied at scale in much smaller human embryonic tissues, because we integrated data from many tissues we could correlate enhancer activity to target genes over megabase distances (Figure 7). Where correlations are linked to expression of the same gene, it then becomes possible to group enhancers. Alongside the need for larger patient cohorts, grouping individual enhancers into larger clusters should increase statistical power, which can be otherwise limiting when causally linking non-coding elements to developmental disorders.

Deciphering profiles of H3K27me3 alongside other regulatory marks and expression profiles was also informative. We did not observe bivalent marking of developmental promoters ‘poised’ for gene expression. Instead, we discovered that organ-specific developmental programmes were disallowed in other human embryonic tissues by active repression at a series of gene promoters. The ontology of these gene sets, including many encoding TFs, inferred they are an important aspect of ensuring correct cell fate decision. This realization opens up a new opportunity for more rigorous benchmarking of differentiated hPSCs, including organoids, both for proximity to the intended lineage in how appropriate gene expression is activated but also against a clearly defined set of epigenomic features for how undesired cell fates are avoided.

In summary, we present an integrated atlas of epigenomic regulation and transcription responsible for human organogenesis and make all datasets freely available. The uncovering of novel regulatory regions and patterns of regulation across organs arose because of direct study of human embryonic tissue. The data complement current international projects such as the Human Cell Atlas^34^, by providing greater resolution of regulatory information and depth of sequence information. Moreover, our integrated analyses establish a new framework for prioritising and interpreting disease-associated variants discovered by WGS^35^ and provide clear routes towards understanding the underlying mechanisms.

## MATERIALS & METHODS

### Sample dissection

Human embryonic material was collected under ethical approval, informed consent and according to the Codes of Practice of the Human Tissue Authority as described previously^15^. Tissue collection took place on our co-located clinical academic campus overseen by our research team ensuring immediate transfer to the laboratory. Material was staged by the Carnegie classification and individual tissues and organs were immediately dissected (Supplementary table 1). The material collected here for epigenomic analysis was matched to material isolated for a previous transcriptomic study^15^ and the dissection process was identical. In brief, the pancreas, adrenal gland, whole brain, heart, kidney, liver, limb buds, lung, stomach, and anterior two-thirds of the tongue were visible as discrete organs and tissues. All visible adherent mesenchyme, including capsular material (adrenal), was removed under a dissecting microscope. The ureter was removed from the renal pelvis. A window of tissue was removed from the lateral wall of the left ventricle of the heart. The dissected segment of liver avoided the developing gall bladder. The trachea was removed where it entered the lung parenchyma. The stomach was isolated between the gastro-oesophageal and pyloric junctions. The palatal shelves were dissected on either side of the midline. The eye was dissected and the RPE peeled off mechanically from its posterior surface (facilitated by the dark pigmentation of the RPE allowing straightforward visualisation).

Tissues were gently teased apart before cross-linking in 1% formaldehyde for 10 min at room temperature. Fixation was quenched with 125 mM glycine for 5 min at room temperature before centrifugation, removal of the supernatant and washing twice with 1 ml PBS. The final PBS supernatant was discarded and samples stored at −80 °C until use (Supplementary table 1).

### Chromatin immunoprecipitation (ChIP), RNA isolation and sequencing

All ChIPseq datasets were in biological replicate except for stomach and tongue (Supplementary table 1). Each sample was placed in lysis buffer [10 mM HEPES, 0.5 mM EGTA, 10 mM EDTA, 0.25% Triton X-100 and protease inhibitor cocktail (Roche)] on ice for 5 min and nuclei released with 10 strokes in a Dounce homogeniser. Nuclei were pelleted by centrifugation at 700 rcf for 10 min at 4 °C and the supernatant discarded. Nuclei were resuspended in ice cold wash buffer (10 mM HEPES, 0.5mM EGTA, 1 mM EDTA, 20 mM NaCl and protease inhibitor cocktail) then pelleted by centrifugation at 700 rcf for 10 min at 4 °C and the supernatant discarded. Nuclei were lysed (50 mM Tris HCl, 10 mM EDTA, 1% SDS and protease inhibitor cocktail) and sonicated under prior optimised conditions (Diagenode Bioruptor). Sufficient sample was prepared to allow in parallel immunoprecipitation for H3K4me3, H3K27ac and H3K27me3 to minimise technical variation. 1 μg DNA equivalent was used for each pulldown. Samples were diluted with 9 volumes of dilution buffer (16.7 mM Tris-HCL, 1.2 mM EDTA, 167 mM NaCl, 0.01% SDS and 1.1% Triton X-100). 20 μl ChIP grade magnetic beads were washed twice in dilution buffer and incubated with each sample for 3 h on a tube rotator at 4 °C to preclear the sample. The beads were separated and the pre-cleared lysate transferred to a separate tube. The magnetic bead pellet was discarded. For each histone modification 3 μg of antibody (Supplementary table 1) were added to each sample followed by incubation on a tube rotator at 4 °C overnight. 30 μl magnetic beads were washed twice in immunoprecipitation dilution buffer and incubated with samples for 3 h at 4 °C. Beads were collected and washed twice with wash buffer A (20 mM Tris-HCl, 2 mM EDTA, 50 mM NaCl, 0.1% SDS and 1% Triton X-100), once with wash buffer B (10 mM Tris-HCl, 1 mM EDTA, 250 mM LiCl, 1% NP40 and 1% Deoxycholate) and twice with TE buffer (10 mM Tris-HCl and 1 mM EDTA). Beads were then incubated in elution buffer (1% SDS and 100 mM NaHCO3) for 30 min at 65 °C and the beads discarded. The resulting samples were incubated with 167 mM NaCl for 5 h at 65 °C to remove crosslinks followed by 1 h incubation with 14 μg proteinase K. The resulting chromatin was then purified (MinElute, QIAGEN).

DNA libraries were constructed according to the TruSeq^®^ ChIP Sample Preparation Guide (Illumina, Inc.). Briefly, sample DNA (5-10 ng) was blunt-ended and phosphorylated, and a single ‘A’ nucleotide added to the 3’ ends of the fragments in preparation for ligation to an adapter with a single base ‘T’ overhang. Omitting the size selection step, the ligation products were then PCR-amplified to enrich for fragments with adapters on both ends. The final purified product was then quantitated prior to cluster generation on a cBot instrument (Illumina). The loaded flow-cell was sequenced (paired-end) on a HiSeq2500 (Illumina). In total, ChIPseq was carried out in three batches with hierarchical clustering analysis to examine for batch effect (Supplementary figure 7).

RNAseq for this study has been described previously^15^; using identical methodology, we added single datasets for pancreas and tongue and two datasets for lung to create biological transcriptomic replicates for all tissues (Supplementary table 2).

### Mapping of ChIPseq and RNAseq

The first batch of ChIPseq was mapped originally to hg19 using Bowtie 1.0.0 (parameters -m1 - n2 -l28, uniquely mapped reads only)^36^ and peaks called using MACS2 (2.0.10.20131216)^37^ against a common input sample (derived from all tissues). To prioritise candidate enhancers for transgenic testing, H3K27ac data from ENCODE (7 cell lines) and NIH Roadmap (154 samples)^10,23^ were mapped similarly. Subsequently, all data, including the external H1 hPSC and adult pancreas data (Figure 3c), were mapped to hg38 using STAR (2.4.2a)^38^. ChIPseq reads were trimmed to 50 bp for consistency and only uniquely mapped reads were retained. GENCODE 25 gene annotations were used for RNAseq mapping and read counting^39^.

### Chromatin and promoter state analysis

Genomic segmentation was performed using chromHMM (version 1.11)^17^ labelling samples by tissue and histone modification. The three histone marks allowed for eight segment states.

Clustered promoter states were identified for an annotated set of 19,791 protein-coding genes in each tissue using ngs.plot on unnormalized reads for the combined dataset of replicated RNAseq and ChIPseq for H3K4me3, H3K27ac and H3K27me3^21^. Default settings allowed for five clusters based on rank profiles of read counts 3 kb either side of the TSS. The returned clusters were then classified according to characteristics detected in both replicates: ‘Actively repressed’ (H3K27me3 signal >50% of maximum and mean transcript counts <10% of maximum); ‘Narrow expressed’ [H3K4me3 signal >25% of maximum with > 90% of reads downstream of the TSS and skew >0.65 (measured across 100 equidistant percentiles from TSS to +3 kb); and mean transcript counts >10% of maximum]; ‘Broad expressed’ (as for ‘Narrow expressed’ but with skew <0.65); ‘Bi-directional expressed’ (H3K4me3 signal >25% of maximum with <90% of reads downstream of the TSS; and mean transcript counts >10% of maximum); ‘Bi-dir2’ (as for ‘Bi-directional expressed’ but without the H3K4me3 signal); ‘Expressed2’ (H3K4me3 signal >25% of maximum with mean transcript counts <10% of maximum); and ‘Inactive’ (<25 of maximum for H3K4me3 and H3K27me3 and mean transcript counts <10% of maximum). This approach left each gene uniquely assigned to one cluster in any tissue. ‘Bi-dir2’ was only identified in RPE (Supplementary figure 2). ‘Expressed2’ was detected in lung, liver and brain (Supplementary figure 3). While superficially this category lacked significant transcription, in fact, total gene-level read counts were very similar to ‘Broad expressed’. However, longer mRNA and longer first introns limited transcript detection at the TSS (Supplementary figure 4). The full listings are in Supplementary table 3.

The over-representation of TFs in the TSS regions marked with H3K27me3 and featuring CpG islands was assessed on the dataset of 1,659 genes encoding all the TFs compared against the remaining 18,132 non-TF genes using Fisher’s exact test (two-sided).

Alluvial plots were created using the R package Alluvial Diagrams version 0.2-0^40^ with modification of the R code to reorder the horizontal splines (alluvia) within each tissue to keep similar colours together.

### Transgenic analysis in zebrafish

A systematic approach identified candidate enhancers that were human embryo-enriched and tissue-specific. We identified marks from the first batch of H3K27ac with RPKM ≥ 25 and ≥ 2.5-fold enrichment in the human embryo compared to ENCODE (7 cell lines)^23^ or NIH Roadmap datasets (154 tissues, including fetal datasets from the second trimester)^10^; and that were undetected in the FANTOM5 project^24^. To filter these embryonic marks for tissue specificity an initial dataset was selected at random and peaks called that were > 200bp. The H3K27ac datasets from other embryonic tissues were then overlaid sequentially in random order. Only called peaks > 200 bp were included. After each addition, any peaks with < 50% overlap between the new and existing dataset were retained. For those retained regions overlapping sequence was filtered out. Once completed, the final set of human embryo-enriched, tissue-specific sequences were again filtered for regions >200 bp. Re-running the tissue specificity algorithm for random addition of datasets resulted in a 99.6% match to the first analysis. These candidate enhancer regions were filtered for sequence conservation (PhastCons LOD score >50)^41^ and correlated with surrounding transcription (≤1 mb in either direction). We manually inspected the remainder for proximity (<1 mb) to genes encoding TFs associated with major developmental disorders and ensured no H3K4me3 or polyadenylated transcription in the immediate vicinity (i.e. an unannotated promoter). This resulted in 44 candidate enhancers from which we tested ten. The candidate sequences were first cloned in TOPO vector using pCR8/GW/TOPO TA cloning kit (Catalogue number K252020, Invitrogen Thermo Fisher Scientific) and then recombined to the reporter vector Minitol2-GwB-zgata2-GFP-48^42^ using the Gateway LR clonase II Enzyme mix (Cat. No. 11791020, Invitrogen Thermo Fisher Scientific). The reporter vector contains a robust midbrain enhancer as an internal control for transgenesis.

Transgenic fish were generated with the Tol2 transposon/transposase method of transgenesis^43^. *Danio rerio* embryos were collected from natural spawning and injected in the yolk at the one-cell stage. The injection mixture contained 50 ng/μl Tol2 transposase mRNA, purified enhancer test vector and 0.05% phenol red. The concentration of the enhancer test vector was between 15 and 30 ng/μl. Injected embryos were visualized from 24 hpf to 48 hpf in an Olympus stereomicroscope coupled to a fluorescence excitation light source in order to detect the pattern of GFP.

Embryos and adults zebrafish were maintained under standard laboratory conditions. They were manipulated according to Spanish and European regulation. All protocols used have been approved by the Ethics Committee of the Andalusian Government (license numbers 450-1839 and 182-41106 for CABD-CSIC-UPO).

### Genome binning, normalisation and thresholding

The genome was parsed into 3,087,584 non-overlapping contiguous 1 kb bins to compare ChIPseq profiles across tissues and replicates. Reads were counted into bins according to their mapped start position using csaw^44^. Reads from mitochondrial and unplaced chromosome annotations were removed. A further 697 bins were filtered out for possessing >10,000 reads in all samples or if the mean read count from input controls was ≥50% of the mean read count of all samples or for being situated in pericentromeric regions (using table ideogram from UCSC; listed in Supplementary table 4). For correlations with surrounding transcription binned read counts were down-sampled statistically using subSeq^45^ weighting each sample by the value of the 99th percentile.

Downsampling of read counts to the 99^th^ percentile was used to generate the custom ‘elbow’ threshold that called bins as marked or not for subsequent downstream analyses. When read counts were ordered and plotted by rank the resulting graph was typically exponential with most bins having zero or very few reads (below the elbow threshold) and a small number of bins with very high read counts (above the elbow threshold). The elbow was defined as the point on the line with the shortest Euclidian distance to the maximum rank intercept with the x-axis. Our code (arseFromElbow) to determine these thresholds from a vector of counts is available on github^46^. Phi correlation was used to measure the agreement between tissue replicates called by the 1 kb binning method compared to MACS^37^. Hierarchical clustering of datasets was undertaken to assess potential batch effect (displayed by heatmap) based on the combined set of the 10,000 most highly ranked bins from each sample. Sets of tissue-specific (replicated in exactly one tissue) and tissue-selective bins (replicated in a given tissue and up to a half of all samples) were produced for each embryonic tissue. EulerGrids showing pattern frequencies of bins across samples were produced using the function plotEuler^47^ as an adaptation of a proposal from Reynolds and colleagues^48^ on Biostars.org^49^.

### Annotation set enrichment for genes and genomic regions

Lists of genes and genomic regions (e.g. 1 kb bins) were tested for enrichment of annotations using the R package XGR version 1.1.1 under default parameters^50^. For TSS clusters (e.g. Alluvial plots) only the remaining annotations used in the ngs.plots were included as background.

### Motif analysis

HOMER v4.9 was used to search for enriched motifs in selected sets of bins^51^. For selected 1 kb bins marked with H3K27Ac, the background set was the remainder of bins with replicated H3K27Ac across all tissues [n=160,043].

## ACKNOWLEDGEMENTS

We are very grateful to all women who consented to take part in our research programme and for the assistance of research nurses and clinical colleagues at the Manchester University NHS Foundation Trust. We thank Peter Briggs and Andy Hayes of the Bioinformatics and Genomic Technologies Core Facilities at the University of Manchester. The work was supported by Wellcome grants 088566, 097820 and 105610, with additional support from MRC project grants MR/L009986/1 to NB and NAH, MR/J003352/1 to KPH, and MR/000638/1 and MR/S036121/1 to NAH. REJ was an MRC clinical research training fellow and SJW was an MRC doctoral account PhD student. JLGS was supported by the Marató TV3 Fundation (Grant 201611).

## AUTHOR CONTRIBUTIONS

DTG, AAB and NAH devised the study and planned experiments. KPH, MB, SJW, REJ, ADS and NB were involved in study design and oversight of human embryonic material collection (REJ, KPH). AAB processed the human embryonic material and prepared samples for all sequencing analyses. DTG and ID conducted the bioinformatics analyses. JLGS, SJG, PNF undertook the transgenic analyses. NAH, DTG, PS and MEH conducted the analysis of developmental disorders. DTG and NAH wrote the manuscript with input from AAB and editing from KPH, NB, ADS and JLGS. NAH is the guarantor.

## AUTHOR INFORMATION

Novel ChIPseq and RNAseq reads have been deposited in the European Genome Phenome repository under accessions: EGAS00001003738 and EGAS00001003163.

To view data in the UCSC genome browser, a trackhub is available at http://www.humandevelopmentalbiology.manchester.ac.uk/. The authors declare no competing financial interests.

## SUPPLEMENTARY TABLES

Five supplementary tables are available in the appended file (.xls). The legends for individual supplementary tables are contained within the file worksheets.

## SUPPLEMENTARY FIGURES

**Supplementary figure 1 (relates to Figure 1).**
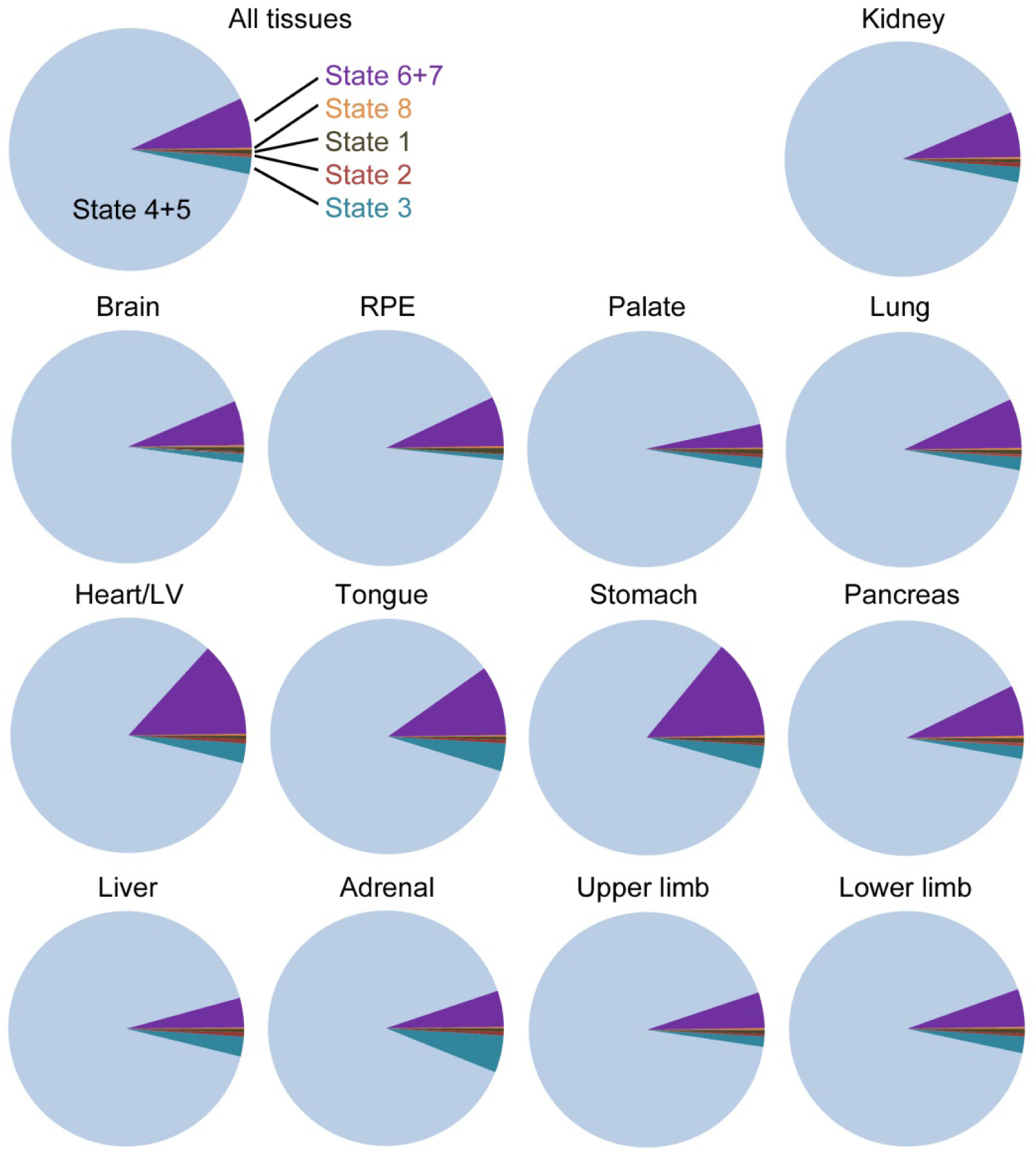
Genome coverage for the different histone modifications in all tissues. Pie charts for individual tissues of genome coverage according to chromatin state by ChromHMM. The average for all tissues is shown in Figure 1c.

**Supplementary figure 2 (relates to Figure 2).**
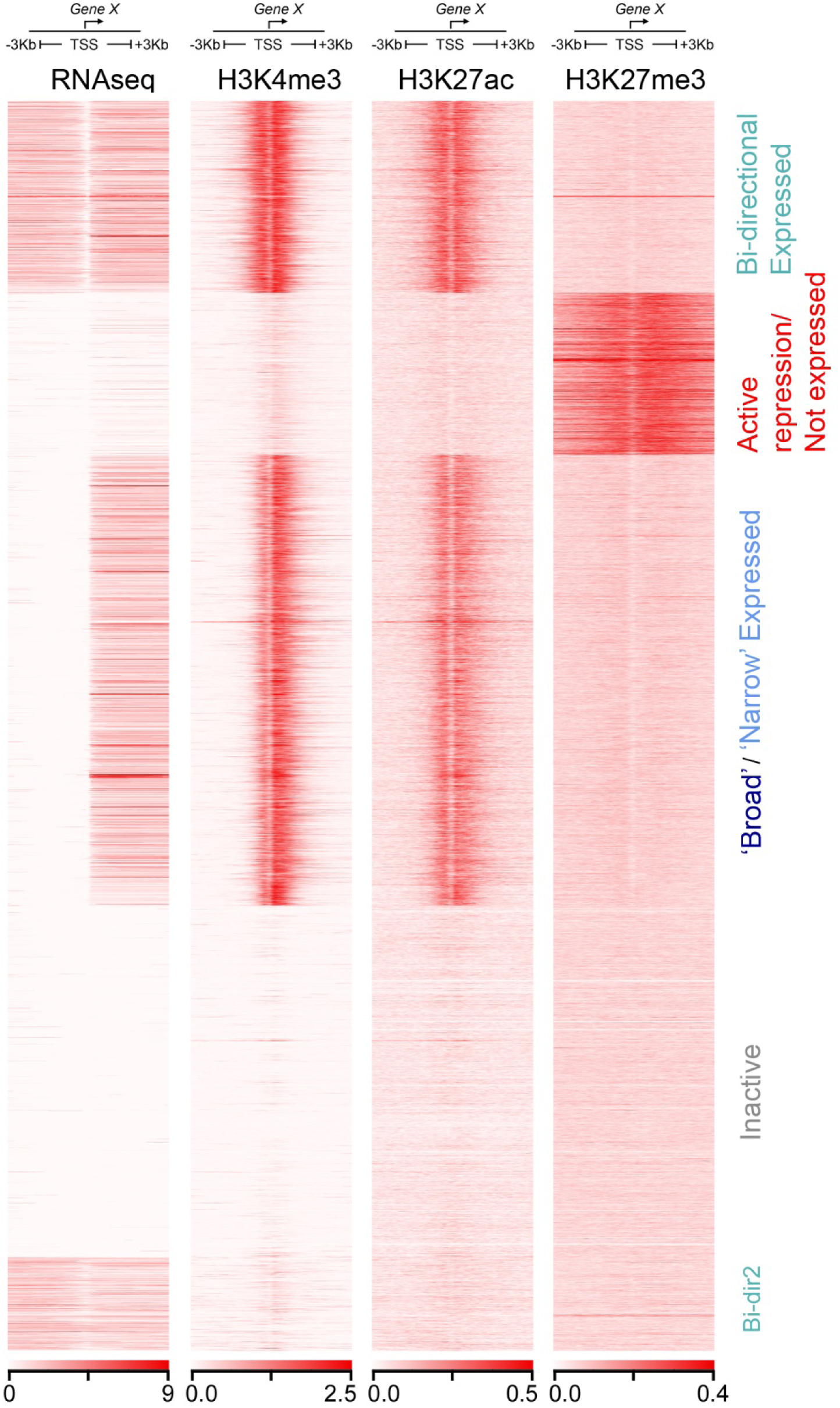
Variant promoter state observed for retinal pigmented epithelium (RPE). In RPE the ‘broad’ and ‘narrow’ expressed categories identified in most tissues and shown in Figure 2a were aggregated by ngsplot and termed ‘broad/narrow’ expressed. This occurred in favour and as a consequence of clustering a subset of genes with less robust bidirectional transcription and barely any marking for H3K4me3 or H3K27ac (termed ‘Bi-dir2’).

**Supplementary figure 3 (relates to Figure 2).**
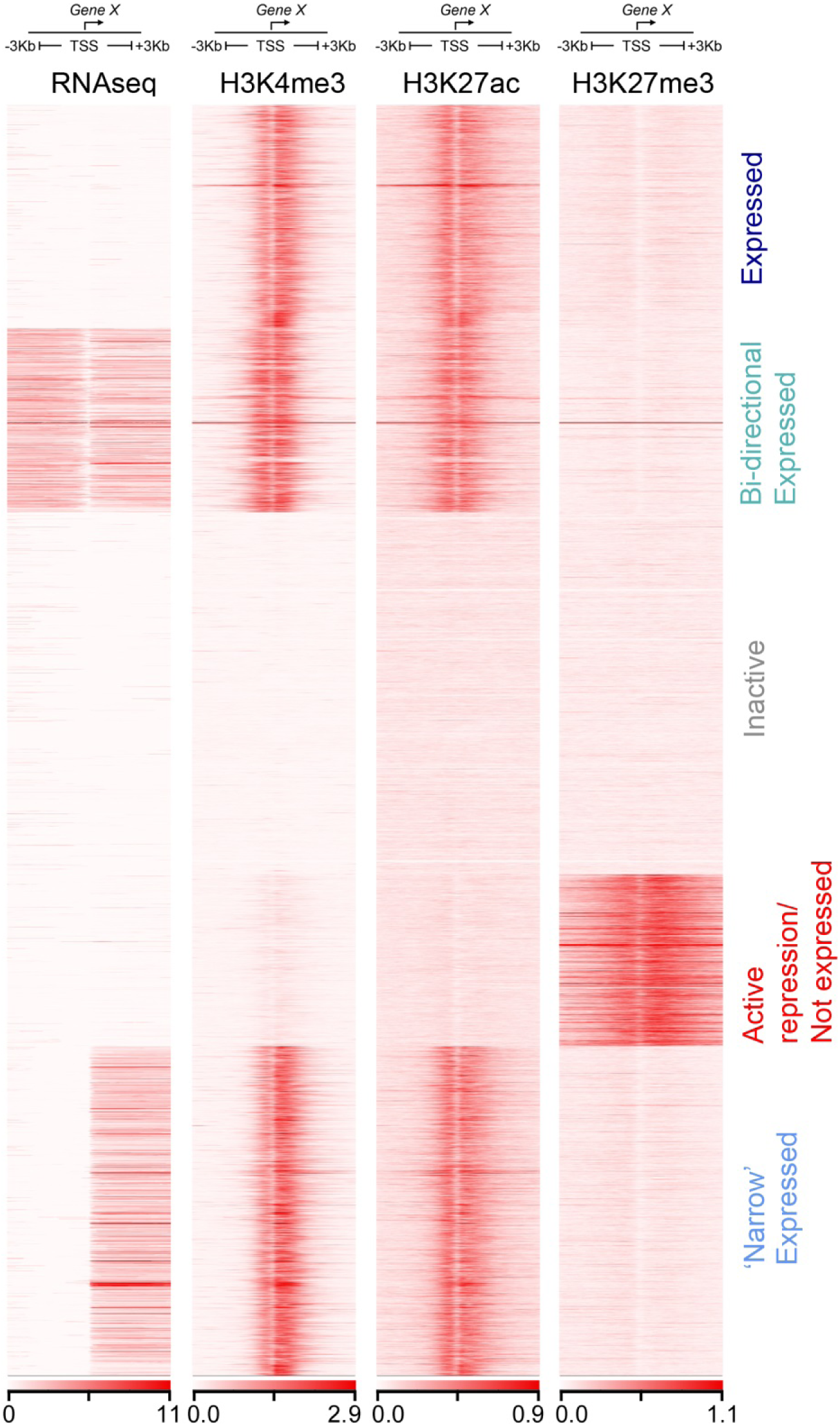
Variant promoter states observed for brain, lung and liver. In brain, lung and liver the ‘Broad expressed’ category, evident in Figure 2a, is marked as ‘Expressed’ (example shown for lung). While the H3K4me3 and H3K27ac marks were indistinguishable from the ‘Broad expressed’ category of Figure 2a, there was no accompanying RNAseq detection at the TSS. These genes encoded longer transcripts with characteristically long first introns (Supplementary figure 4) leading to an under-representation of RNAseq reads at the TSS. Total transcript count across the entire gene was equivalent for ‘Broad expressed’ (Figure 2a) and ‘Expressed’ genes.

**Supplementary figure 4 (relates to Figure 2 and Supplementary figure 3).**
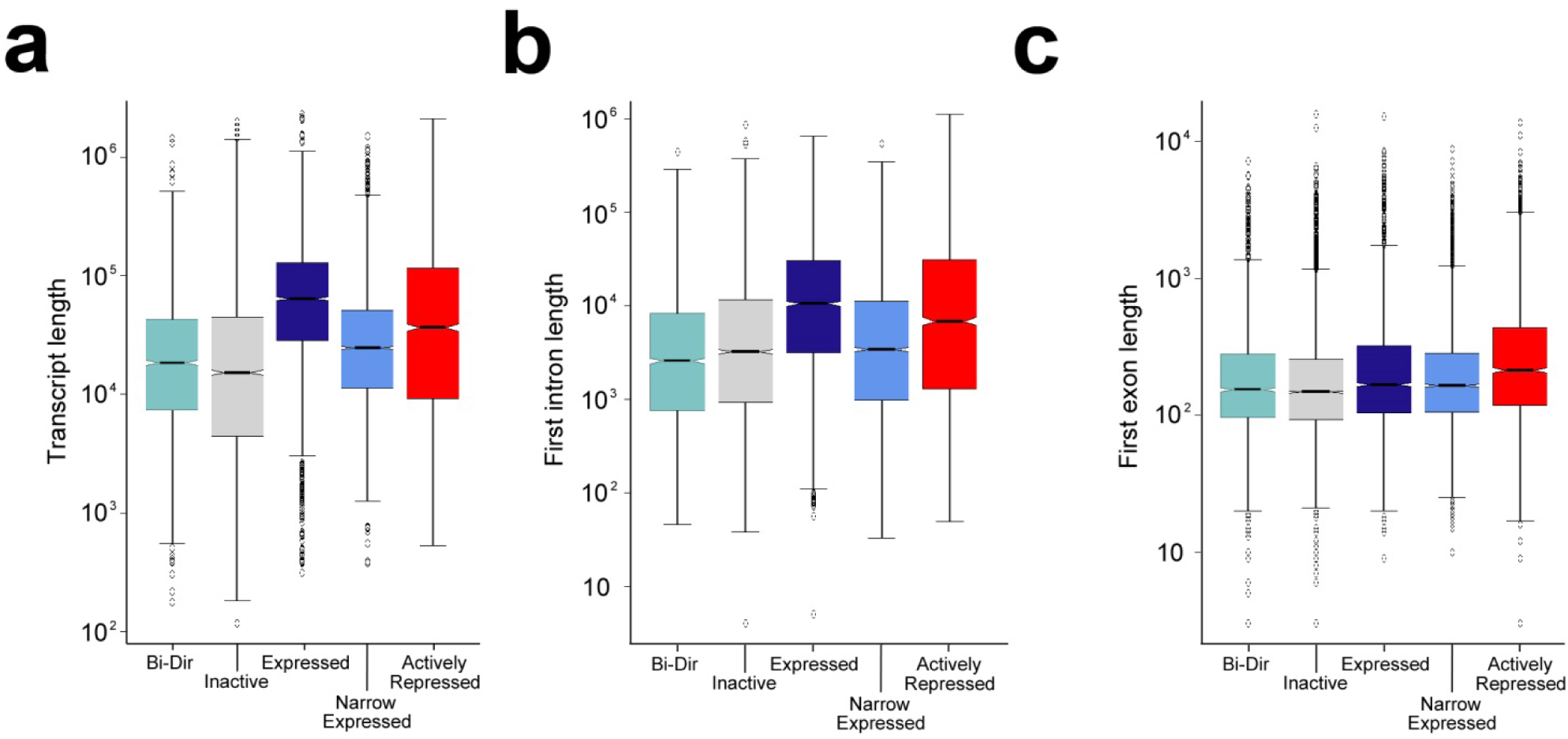
Characteristics of the ‘Expressed’ promoter state (brain, lung and liver). a) Transcript length b) first intron length and c) first exon length for the categories of genes detected in brain, lung and liver (Supplementary figure 3). Although overall transcript counts for ‘Expressed’ were similar to ‘Broad Expressed’ (Figure 2a), RNAseq was not detected at the TSS due to longer first introns and overall transcript length. The first exon was similar to other categories. The example shown is for lung with the same findings observed for brain and liver.

**Supplementary figure 5 (relates to Figure 2).**
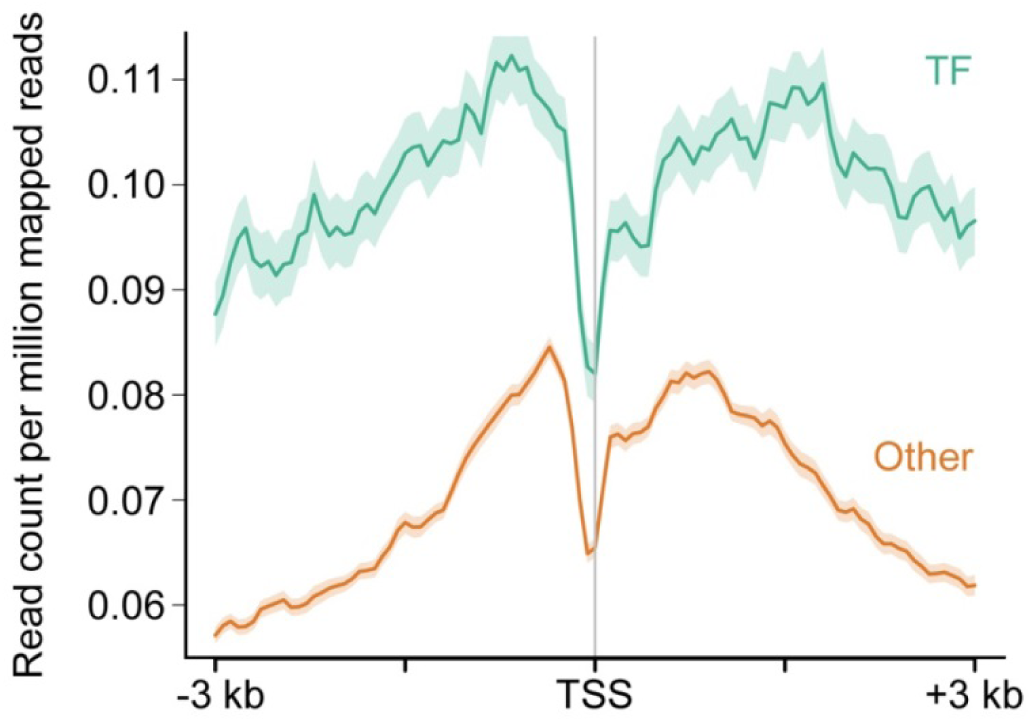
Levels of H3K27me3 at the TSS of actively repressed genes which encoded transcription factors (TF) compared to those encoding all other proteins (Other). Within the ‘Active Repression’ category across all tissues genes encoding transcription factors (TFs) possessed appreciably greater marking with H3K27me3 at their transcriptional start site (TSS) compared to those genes encoding other proteins.

**Supplementary figure 6 (relates to Figure 3).**
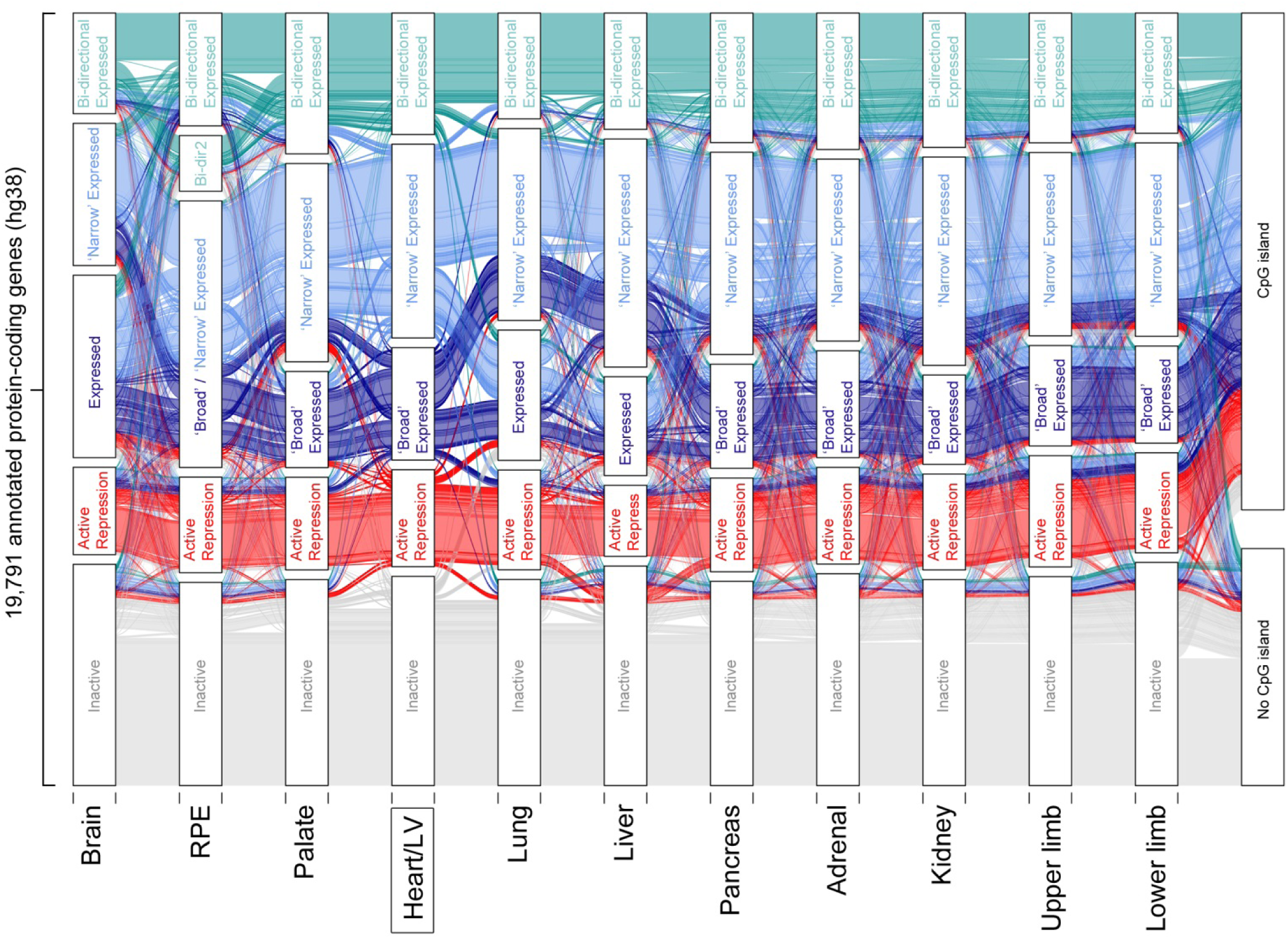
Integration of all identified promoter states across tissues. Alluvial plot showing promoter state for 19,791 annotated genes across all tissues with replicated datasets. All the amalgamated promoter states associated with gene transcription in Figure 3 are shown individually here: ‘Broad expressed’, ‘Narrow expressed’, ‘Expressed’, ‘Bidir’ and ‘Bidir2’. The plot shown is centred on (and with variance from) the promoter state in the Heart/LV dataset. The plot also categorises genes according to the presence or absence of a CpG island at the promoter. Genes with an ‘Inactive’ promoter state characteristically lacked a CpG island. Promoters either actively transcribed or repressed tended to possess a CpG island.

**Supplementary figure 7.**
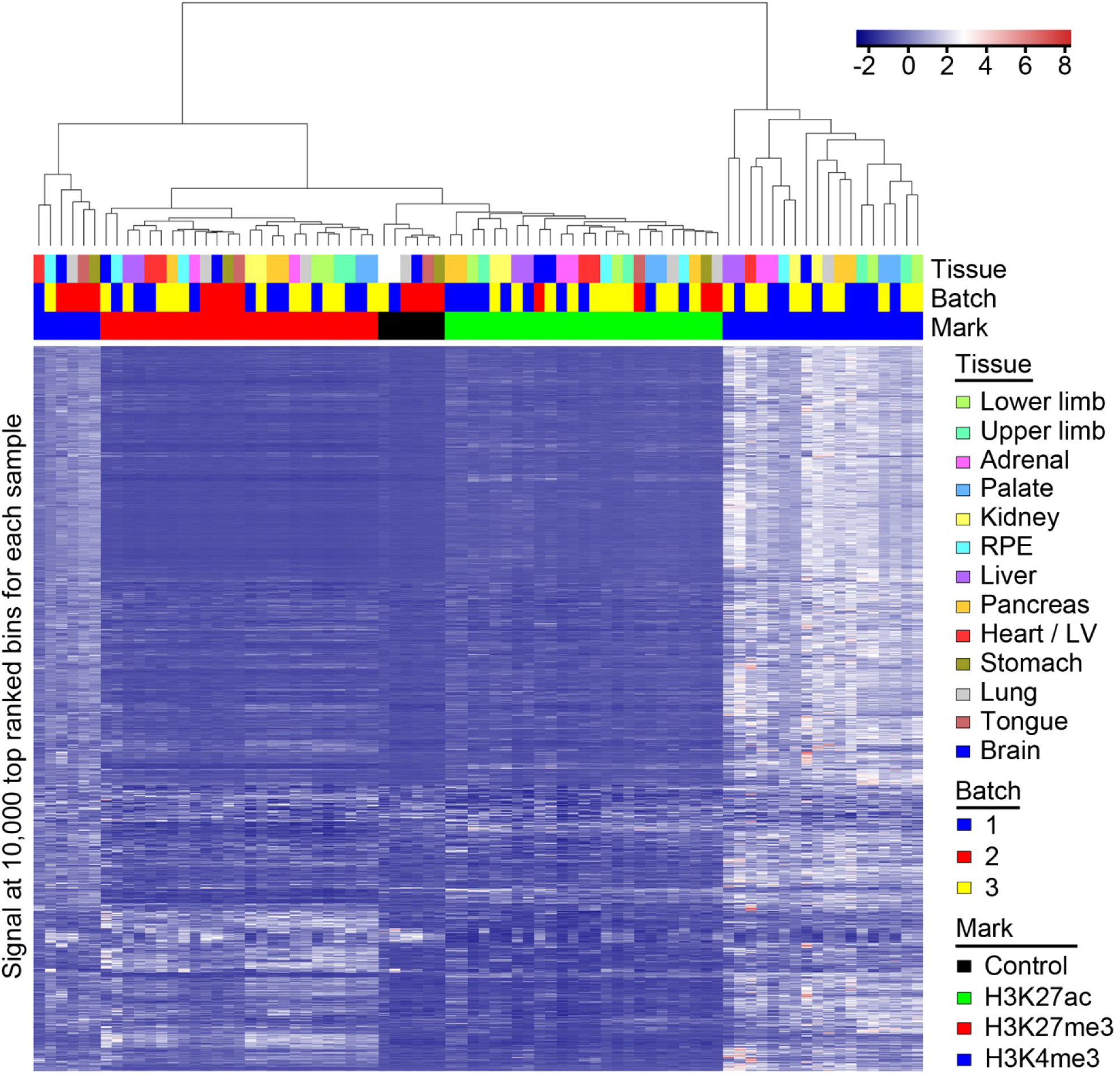
Heatmap showing hierarchical clustering of ChIPseq datasets based on the combined set of 10,000 most highly ranked bins from each sample. Samples clustered according to mark and did not cluster according to sequencing batch.

**Supplementary figure 8.**
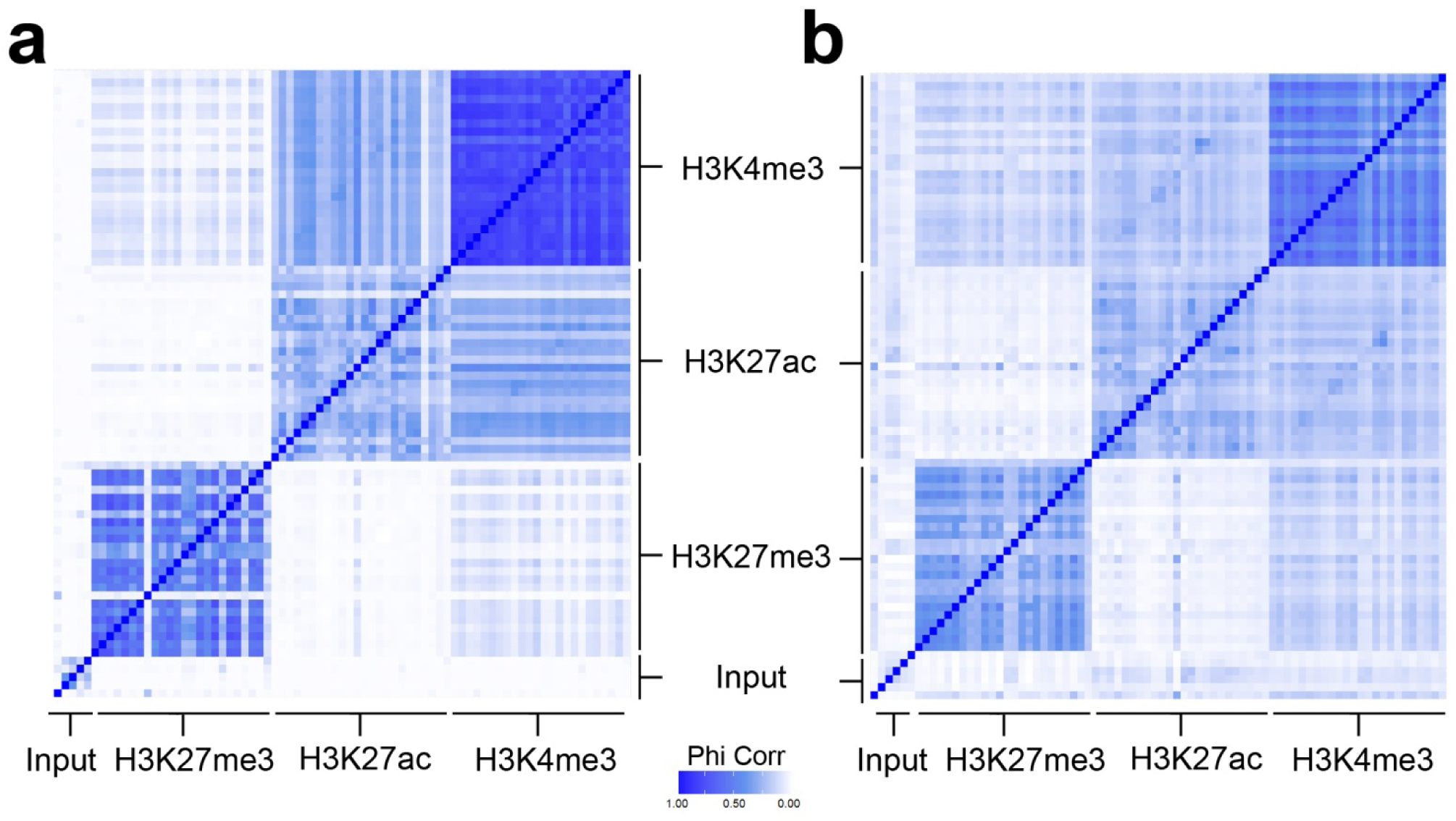
Heatmap showing Phi correlation between samples when peaks are called by MACs or allocated according to read count into 1 kb bins. a) Peak calling by MACs^37^. b) Allocation according to read counts into 1 kb bins. The two approaches produced similar heatmaps thus benchmarking the 1 kb bin approach. Each histone modification is comprised of 26 individual rows and columns for the 12 tissue replicates plus single datasets for tongue and stomach. The input is comprised of 5 control samples. The dark blue diagonal line is the perfect correlation from assessing each sample against itself.

**Supplementary figure 9.**
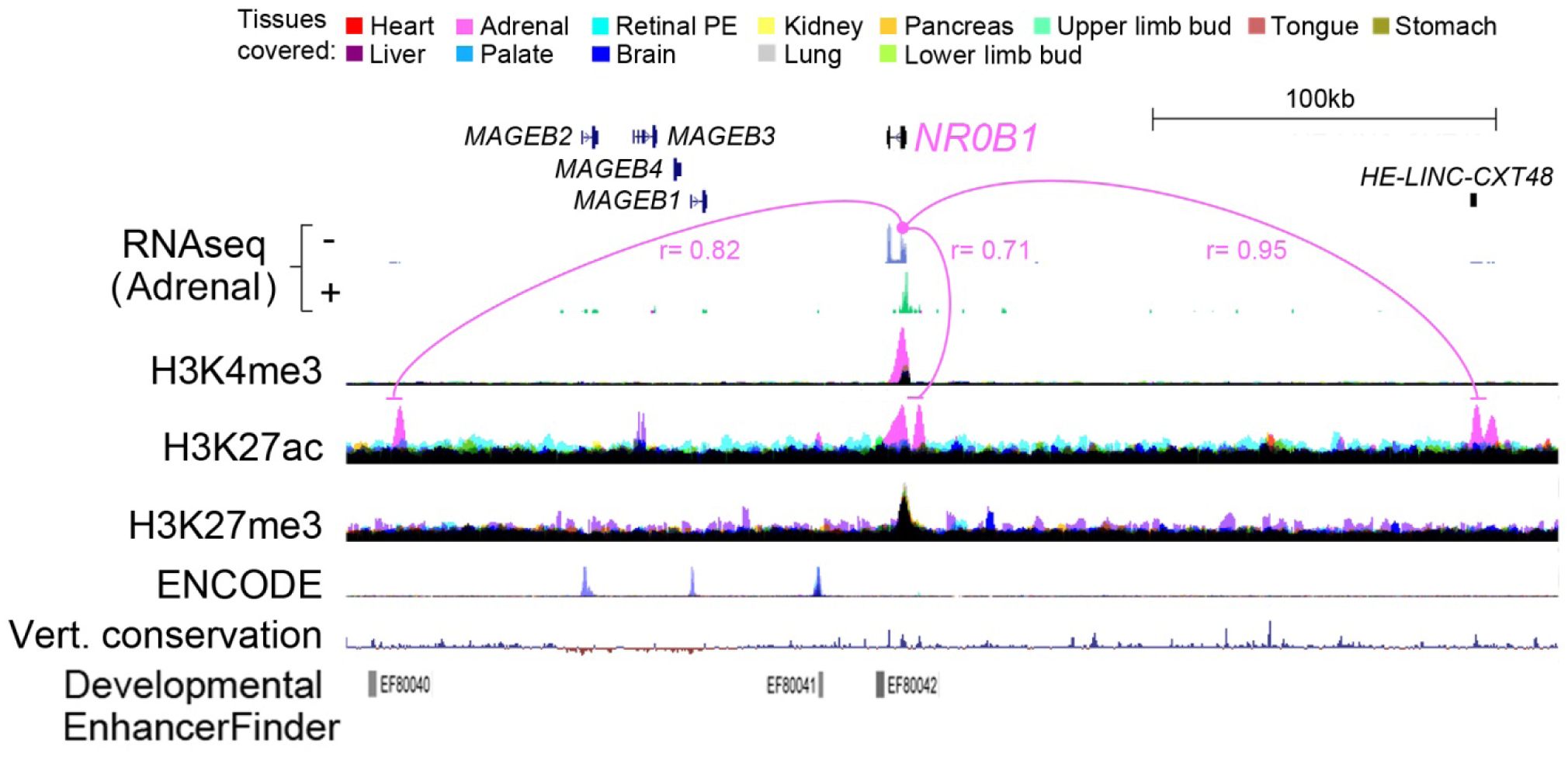
Adrenal-specific epigenomic landscape over 300 kb at the *NR0B1* locus on the X chromosome. Assembled tracks show RNAseq for the adrenal and multi-layered data for all tissues for each histone modification. One replicate of each track is shown for simplicity. *NR0B1* was only expressed in the adrenal and has an adrenal-specific H3K4me3 mark. Multiple adrenal-specific H3K27ac enhancer peaks were visible across 300 kb, all highly correlated with the expression of *NR0B1*. Some of the enhancers are poorly conserved, unpredicted by in silico tools (Developmental Enhancer Finder^53^), and absent from ENCODE datasets^23^. Identifying multiple enhancers facilitates their grouping to assist with statistical power when assessing the potential pathogenicity of patient variants in whole genome sequencing data.

**Supplementary figure 10 (relates to Figure 5).**
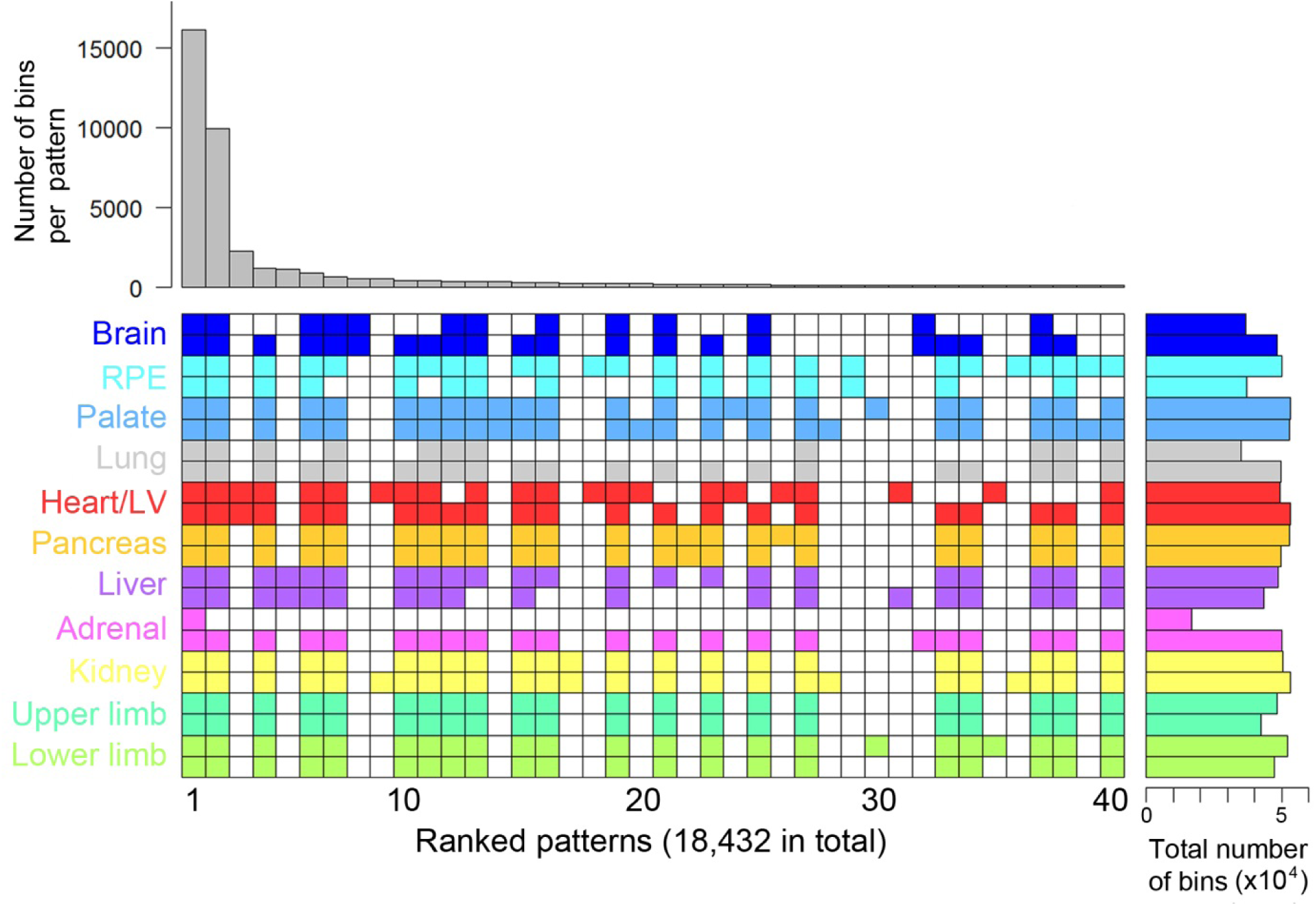
Patterns of H3K4me3 across human embryonic tissues with the requirement of detection in at least 2 samples. Euler grid for bins marked by H3K4me3 (defined by elbow plots) in replicated tissues (i.e. two rows/replicates per tissue). Total number of marked bins per individual dataset is shown to the right. The grid shown required a bin to be called in any two or more samples and is ordered by decreasing bin count per pattern (bar chart above the grid). A total of 18,432 different patterns were identified (far fewer that the 48,570 found for the corresponding analysis of H3K27ac). The top 40 are shown. All tissue-specific patterns emerged in the top 2,267 (within the top 12.3% of patterns).

**Supplementary figure 11 (relates to Figure 5).**
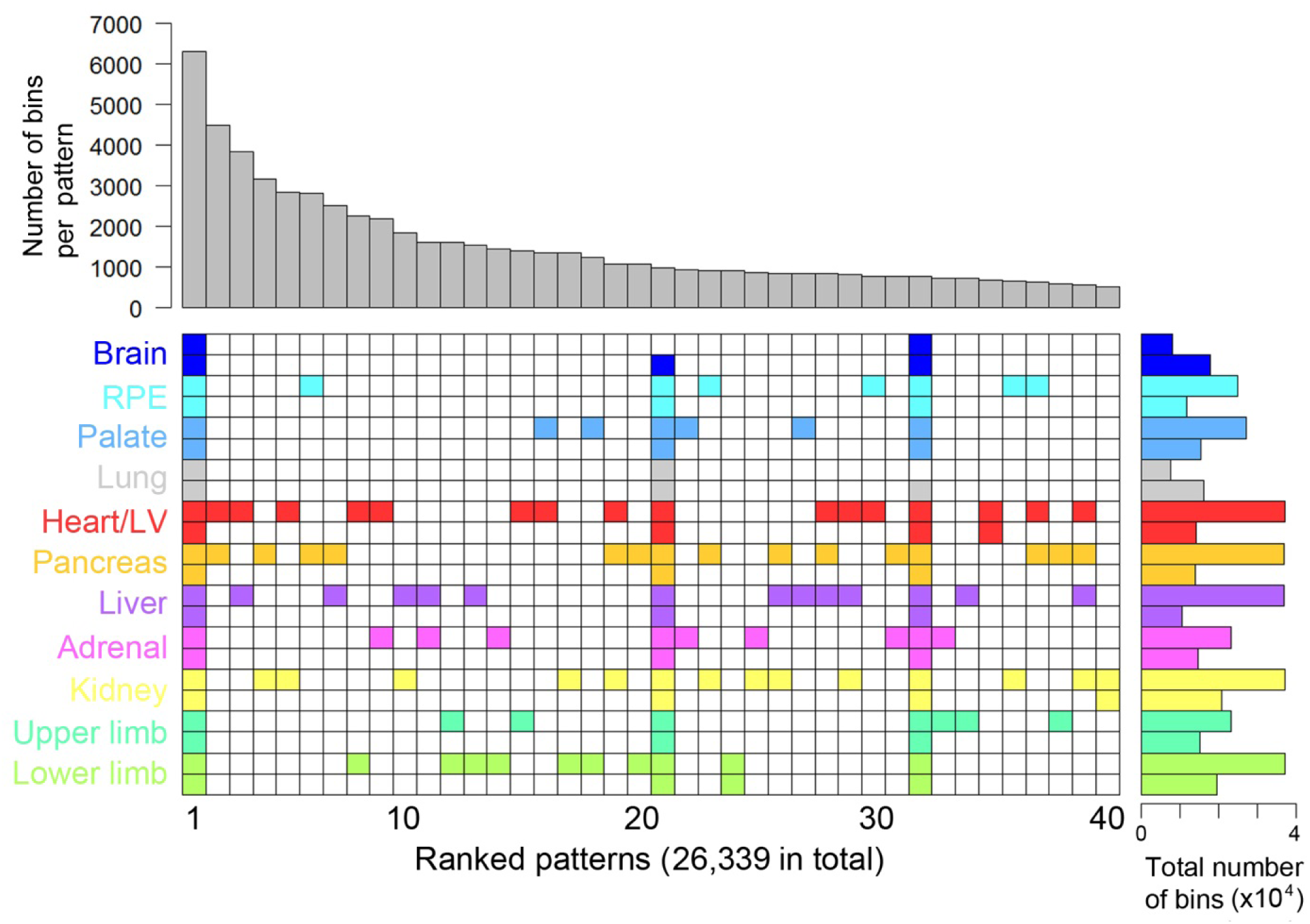
Patterns of H3K27me3 across human embryonic tissues with the requirement of detection in at least 2 samples. Euler grid for bins marked by H3K27me3 (defined by elbow plots) in replicated tissues (i.e. two rows/replicates per tissue). Total number of marked bins per individual dataset is shown to the right. The grid shown required a bin to be called in any two or more samples and is ordered by decreasing bin count per pattern (bar chart above the grid). A total of 26,339 different patterns were identified (far fewer that the 48,570 found for the corresponding analysis of H3K27ac). The top 40 are shown. All tissue-specific patterns emerged in the top 836 (within the top 3.2% of patterns).

**Supplementary figure 12 (relates to Figure 5).**
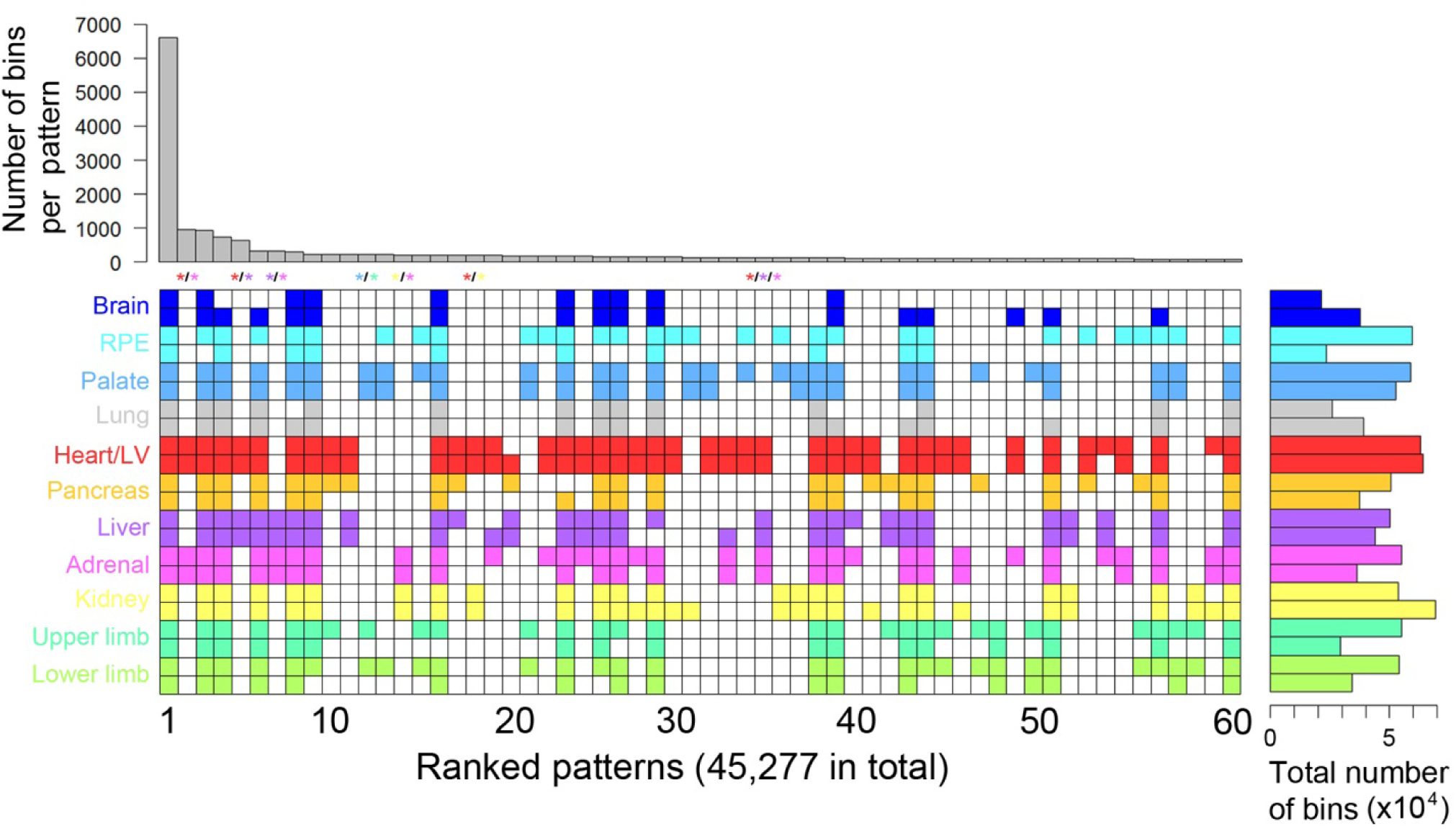
Patterns of H3K27ac across tissues with the requirement of detection in at least 4 samples. Euler grid is shown for bins marked by H3K27ac (defined by elbow plots). Detecting patterns shared across tissues was enforced by detection in at least four samples (which must include at least two tissues). The grid includes replicated tissues (i.e. two rows/replicates per tissue). Total number of marked bins per individual dataset is shown to the right. The grid is ordered by decreasing bin count per pattern (bar chart above the grid). A total of 45,277 different patterns were identified. The top 60 are shown. Colour-coded asterisks above the columns indicate patterns shared uniquely across two (heart-adrenal, heart-liver, palate-limb, kidney-adrenal and heart-kidney) or three (heart-liver-adrenal) tissues.

## REFERENCES

1. Deciphering Developmental Disorders Study: Prevalence and architecture of de novo mutations in developmental disorders. Nature 542, 433–438 (2017).

2. Short, P.J. et al. De novo mutations in regulatory elements in neurodevelopmental disorders. Nature 555, 611–616 (2018).

3. MacArthur, J. et al. The new NHGRI-EBI Catalog of published genome-wide association studies (GWAS Catalog). Nucleic Acids Res 45, D896–D901 (2017).

4. Weedon, M.N. et al. Recessive mutations in a distal PTF1A enhancer cause isolated pancreatic agenesis. Nat Genet 46, 61–64 (2014).

5. van Arensbergen, J. et al. Derepression of Polycomb targets during pancreatic organogenesis allows insulin-producing beta-cells to adopt a neural gene activity program. Genome Res 20, 722–732 (2010).

6. Nord, A.S. et al. Rapid and pervasive changes in genome-wide enhancer usage during mammalian development. Cell 155, 1521–1531 (2013).

7. Schmidt, D. et al. Five-vertebrate ChIP-seq reveals the evolutionary dynamics of transcription factor binding. Science (New York, N.Y.) 328, 1036–1040 (2010).

8. Dickel, D.E. et al. Ultraconserved Enhancers Are Required for Normal Development. Cell 172, 491-499.e415 (2018).

9. Woolfe, A. et al. Highly Conserved Non-Coding Sequences Are Associated with Vertebrate Development. PLOS Biology 3, e7 (2004).

10. Kundaje, A. et al. Integrative analysis of 111 reference human epigenomes (Roadmap Epigenomics Consortium). Nature 518, 317–330 (2015).

11. Cotney, J. et al. The Evolution of Lineage-Specific Regulatory Activities in the Human Embryonic Limb. Cell 154, 185–196 (2013).

12. Wilderman, A., VanOudenhove, J., Kron, J., Noonan, J.P. & Cotney, J. High-Resolution Epigenomic Atlas of Human Embryonic Craniofacial Development. Cell Rep 23, 1581–1597 (2018).

13. Cebola, I. et al. TEAD and YAP regulate the enhancer network of human embryonic pancreatic progenitors. Nat Cell Biol 17, 615–626 (2015).

14. Reilly, S.K. et al. Evolutionary genomics. Evolutionary changes in promoter and enhancer activity during human corticogenesis. Science 347, 1155–1159 (2015).

15. Gerrard, D.T. et al. An integrative transcriptomic atlas of organogenesis in human embryos. eLife 5, e15657.+ (2016).

16. Pollard, K.S., Hubisz, M.J., Rosenbloom, K.R. & Siepel, A. Detection of nonneutral substitution rates on mammalian phylogenies. Genome Res 20, 110–121 (2010).

17. Ernst, J. & Kellis, M. ChromHMM: automating chromatin-state discovery and characterization. Nature Methods 9, 215–216 (2012).

18. Li, F. et al. Bivalent Histone Modifications and Development. Curr Stem Cell Res Ther 13, 83–90 (2018).

19. Harikumar, A. & Meshorer, E. Chromatin remodeling and bivalent histone modifications in embryonic stem cells. EMBO Rep 16, 1609–1619 (2015).

20. Lesch, B.J. & Page, D.C. Poised chromatin in the mammalian germ line. Development 141, 3619–3626 (2014).

21. Shen, L., Shao, N., Liu, X. & Nestler, E. ngs.plot: Quick mining and visualization of next-generation sequencing data by integrating genomic databases. BMC Genomics 15, 284 (2014).

22. Jennings, R.E., Berry, A.A., Strutt, J.P., Gerrard, D.T. & Hanley, N.A. Human pancreas development. Development 142, 3126–3137 (2015).

23. Ernst, J. et al. Mapping and analysis of chromatin state dynamics in nine human cell types. Nature 473, 43–49 (2011).

24. Andersson, R. et al. An atlas of active enhancers across human cell types and tissues. Nature 507, 455–461 (2014).

25. Rana, M.S., Christoffels, V.M. & Moorman, A.F. A molecular and genetic outline of cardiac morphogenesis. Acta Physiol (Oxf) 207, 588–615 (2013).

26. Sizemore, G.M., Pitarresi, J.R., Balakrishnan, S. & Ostrowski, M.C. The ETS family of oncogenic transcription factors in solid tumours. Nat Rev Cancer 17, 337–351 (2017).

27. Martin, H.C. et al. Quantifying the contribution of recessive coding variation to developmental disorders. Science 362, 1161–1164 (2018).

28. Firth, H.V. & Wright, C.F. The Deciphering Developmental Disorders (DDD) study. Dev Med Child Neurol 53, 702–703 (2011).

29. Auton, A. et al. A global reference for human genetic variation. Nature 526, 68–74 (2015).

30. Klopocki, E. et al. Deletions in PITX1 cause a spectrum of lower-limb malformations including mirror-image polydactyly. Eur J Hum Genet 20, 705–708 (2012).

31. Allen, H.L. et al. GATA6 haploinsufficiency causes pancreatic agenesis in humans. Nat Genet 44, 20–22 (2011).

32. Shaw-Smith, C. et al. GATA4 mutations are a cause of neonatal and childhood-onset diabetes. Diabetes 63, 2888–2894 (2014).

33. Won, H. et al. Chromosome conformation elucidates regulatory relationships in developing human brain. Nature 538, 523–527 (2016).

34. Regev, A. et al. The Human Cell Atlas. Elife 6 (2017).

35. Turnbull, C. et al. The 100 000 Genomes Project: bringing whole genome sequencing to the NHS. BMJ 361, k1687 (2018).

36. Langmead, B., Trapnell, C., Pop, M. & Salzberg, S.L. Ultrafast and memory-efficient alignment of short DNA sequences to the human genome. Genome Biology 10, R25 (2009).

37. Zhang, Y. et al. Model-based analysis of ChIP-Seq (MACS). Genome biology 9, R137+ (2008).

38. Dobin, A. et al. STAR: ultrafast universal RNA-seq aligner. Bioinformatics (Oxford, England) 29, 15–21 (2013).

39. Harrow, J. et al. GENCODE: The reference human genome annotation for The ENCODE Project. Genome Research 22, 1760–1774 (2012).

40. Bojanowski, M. & Edwards, R. alluvial: Alluvial Diagrams. https://cran.r-project.org/web/packages/alluvial/index.html (2016).

41. Siepel, A. et al. Evolutionarily conserved elements in vertebrate, insect, worm, and yeast genomes. Genome Research 15, 1034–1050 (2005).

42. Gehrke, A.R. et al. Deep conservation of wrist and digit enhancers in fish. Proc Natl Acad Sci U S A 112, 803–808 (2015).

43. Kawakami, K. et al. A transposon-mediated gene trap approach identifies developmentally regulated genes in zebrafish. Dev Cell 7, 133–144 (2004).

44. Lun, A.T.L. & Smyth, G.K. csaw: a Bioconductor package for differential binding analysis of ChIP-seq data using sliding windows. Nucleic Acids Research 44, e45 (2016).

45. Robinson, D.G. & Storey, J.D. subSeq: Determining Appropriate Sequencing Depth Through Efficient Read Subsampling. Bioinformatics 30, 3424–3426 (2014).

46. Gerrard, D.T. arseFromElbow. utilsGerrardDT/arseFromElbow.R (2019).

47. Gerrard, D.T. plotEuler. https://github.com/davetgerrard/utilsGerrardDT (2019).

48. Reynolds, A. Venn/Euler Diagram Of Four Or More Sets (BioStars.org). https://www.biostars.org/p/77362/#77377 (2013).

49. Parnell, L.D. et al. BioStar: An Online Question & Answer Resource for the Bioinformatics Community. PLOS Computational Biology 7, e1002216 (2011).

50. Fang, H., Knezevic, B., Burnham, K.L. & Knight, J.C. XGR software for enhanced interpretation of genomic summary data, illustrated by application to immunological traits. Genome Med 8, 129 (2016).

51. Heinz, S. et al. Simple combinations of lineage-determining transcription factors prime cis-regulatory elements required for macrophage and B cell identities. Mol Cell 38, 576–589 (2010).

52. http://ms.mcmaster.ca/peter/s743/poissonalpha.html.

53. Erwin, G.D. et al. Integrating diverse datasets improves developmental enhancer prediction. PLoS Comput Biol 10, e1003677 (2014).

